# MECOM promotes leukemia progression and inhibits mast cell differentiation through functional competition with GATA2

**DOI:** 10.1101/2024.05.20.594903

**Authors:** Kohei Iida, Mayuko Nakanishi, Jakushin Nakahara, Shuhei Asada, Tomoya Isobe, Tomohiro Yabushita, Manabu Ozawa, Yasuhiro Yamada, Toshio Kitamura, Keita Yamamoto, Susumu Goyama

## Abstract

MECOM is a nuclear transcription factor essential for the proliferation of hematopoietic stem cells (HSCs) and myeloid leukemia cells. MECOM contains N- and C-terminal zinc finger domains (ZFDs) and binding motifs for the corepressor CtBP to regulate gene expression. Recent studies have shown that germline *MECOM* variants are associated with thrombocytopenia, radioulnar synostosis, and bone marrow failure, collectively termed MECOM-associated syndromes. Although the mutations are clustered in the C-terminal ZFD, how these mutations affect MECOM function has remained unclear. In addition, the individual genes and pathways regulated by MECOM are less well understood. In this study, we showed that the C-terminal ZFD is a major DNA-binding domain of MECOM and that the disease-associated mutations abolish the DNA-binding ability. We also found that MECOM functionally antagonizes GATA2 through the C-terminal ZFD-mediated DNA binding and CtBP interaction, thereby promoting myeloid leukemogenesis while inhibiting mast cell differentiation. Furthermore, we generated mutant MECOM knockin mice harboring a C-terminal ZFD mutation that recapitulate several features of MECOM-associated syndromes, including HSC and B-cell reduction. Our study demonstrates that C-terminal ZFD mutations are loss-of-function mutations with reduced DNA-binding ability, reveals the critical role of MECOM in inhibiting GATA2, and provides a novel mouse model for MECOM-associated syndromes.

## INTRODUCTION

MECOM (<u>M</u>DS1 and <u>E</u>VI1 <u>Com</u>plex Locus) is a member of the PR domain family of transcription factors critical for hematopoietic stem cells (HSCs) [1, 2]. Several studies have shown that loss of MECOM results in the impaired generation and maintenance of HSCs using *Mecom* knockout mice [3,4,5]. On the other hand, aberrant expression of MECOM due to chromosomal translocations, such as inv(3)(q21q26.2) and t(3;3)(q21;q26.2), leads to the onset of myelodysplastic syndrome (MDS) and acute myeloid leukemia (AML) [6,7,8]. Importantly, high MECOM expression is associated with poor prognosis, indicating a potential role for MECOM in maintaining AML stem cells that underlie therapeutic resistance [9,10]. Thus, MECOM is a critical regulator in normal and malignant hematopoiesis.

MECOM contains N- and C-terminal zinc figure domains (ZFD) with distinct DNA-binding specificities [11,12,13]. MECOM has been shown to activate the transcription of several HSC-related genes, including GATA2, PBX1, GFI1 and ERG [5,14,15,16,17]. MECOM has also been shown to repress the expression of several target genes, such as PTEN, through interactions with transcriptional corepressors (C-terminal binding proteins: CtBPs) and epigenetic regulators (SUV39H1, G9a and EZH2) [18,19,20,21,22,23]. In particular, CtBPs (CtBP1 and CtBP2) have been shown to be involved in the MECOM-induced transcriptional repression and leukemogenesis. Despite these advances, the mechanisms of MECOM-mediated transcription and transformation are still not fully understood. For example, it remains unclear which ZFD is more important for MECOM binding to chromatin. Another unresolved issue is the complex relationship between MECOM and GATA2. GATA2 is a versatile transcription factor regulating multiple aspects of hematopoiesis, including HSC expansion, megakaryocyte development, and mast cell development [24,25,26,27,28,29]. We and others have shown that GATA2 is directly upregulated by MECOM, and the MECOM-GATA2 pathway is essential in HSC expansion during embryogenesis [5,30]. On the other hand, it has been shown that the translocations involving the *MECOM* gene at 3q26.2 induced the relocation of the GATA2 enhancer close to the *MECOM* gene, leading to both MECOM overexpression and GATA2 downregulation [31,32]. This clinical observation was also supported by the experiment using a 3q rearranged mouse model showing that GATA2 haploinsufficiency accelerates MECOM-driven leukemogenesis [33,34]. Thus, the role of GATA2 in MECOM-mediated hematopoiesis and leukemogenesis is under debate and appears to be context-dependent.

Recently, heterozygous mutations in *MECOM* genes were identified in patients with Radial Ulnar Synostosis with Amegakaryocytic Thrombocytopenia (RUSAT), an Inherited Bone Marrow Failure Syndrome (IBMFS) [35]. Since patients with the germ line mutations in *MECOM* have shown diverse clinical manifestations, such as bone marrow failure, radioulnar synostosis, B-cell deficiency and hearing loss [36,37,38], the term MECOM-associated syndrome was proposed for this heterogeneous hereditary disease [37]. Importantly, the mutations were concentrated in the C-terminal ZFD of *MECOM* [35,36,37,38,39,40,41], indicating the critical role of the C-terminal ZFD in regulating MECOM function. Indeed, recently generated knockin mice harboring the RUSAT-associated *MECOM* mutation in the C-terminal ZFD exhibited thrombocytopenia and reduced HSCs, which are commonly seen in patients with MECOM-associated syndromes [42]. However, it remained unknown how C-terminal ZFD mutations affect the biological functions of MECOM.

In this study, we performed a detailed functional analysis of MECOM mutants with mutations at the C-terminal ZFD or CtBP-binding sites. These experiments revealed that the C-terminal ZFD functions as the major DNA-binding region of MECOM, while CtBP is required for MECOM-mediated transcriptional repression. We also generated a MECOM mutant knockin mice and confirmed that the C-terminal ZFD mutation (R750W) is indeed a loss-of-function mutation. Interestingly, MECOM has opposing functions to regulate GATA2 expression and function. MECOM binds to the GATA2 promoter through the C-terminal ZFD and activates its transcription. At the same time, MECOM functionally antagonizes GATA2-induced transcription in cooperation with CtBP. This MECOM-mediated functional repression of GATA2 plays an important role in myeloid cell fate specification.

## MATERIALS & METHODS

### Plasmids

Wild-type human MECOM (MECOM-WT) cDNA was obtained from DNAFORM (Clone ID: 100067260, GenBank: BX647613.1) and cloned into pMYs-IRES-GFP (pMYs-IG) vector (Ref) with FLAG tag in N-terminus between EcoRI and NotI sites. MECOM-DL/AS, MECOM-AS/AS, MECOM-R750W and MECOM-C766G mutants were produced by two-step PCR using the FLAG-MECOM-WT as a template. MECOM-WT with 3×HA tag was produced by PCR using the FLAG-MECOM-WT as a template. MECOM-WT and MECOM-R750W with AM tag were produced by PCR using FLAG-MECOM-WT and FLAG-MECOM-R750W as a template, respectively. Human CtBP1 cDNA was obtained from RIKEN BRC DNA Bank (Clone name: IRAL017H06, DDBJ: BC011655), and HA-tagged CtBP1 was cloned into pMYs-IRES-NGFR (pMYs-IN) vector between the EcoR I and Not I sites. FLAG-tagged human GATA2 was cloned into pMYs-IN vector between the EcoR I and Not I sites. To generate the luciferase reporters containing the proximal and distal promoter regions of *GATA2*, the corresponding genomic regions were amplified by PCR from genomic DNA of HEK293T cells. The PCR products were then cloned into pGL4.1 plasmid using Gibson assembly between the Nhe I and Bgl II sites. Primers used are provided in Supplementary Table1.

### Mice

C57BL/6 (Ly5.2) mice (Japan SLC, Inc) were used for colony replating and bone marrow transplantation assays. Rosa26-LSL-Cas9 knockin mice were purchased from Jackson Laboratory (#024857) [43]. To generate EVI1^R751W/WT^ knockin mice, Cas9 protein (IDT, Coralville, IA, USA), gRNA (5’ gggttctcaagtgccgtgtt 3’, IDT) and single-stranded oligodeoxynucleotide (5’ tgctgtggtggaacaacgagctatgctgactttctcttgtggatattttttcttcaaaggtactgtggcaagatatttccaaggtctgcgaaTTt aaca<u>T</u>ggcacttgagaacccacacaggagagcaaccttacaggttagacagtcgtttttctaaatatctcgataaatactataatagtcaatg ctcattaggtca 3’, first two capital Ts or latter underlined single T represent silent mutations to delete PAM or R751W mutation in the KI allele, respectively, IDT) were electroporated into ES cells derived from B6-129F1 mice using Neon Electroporation System (Thermo Fisher, MI, USA). The completion of the desired KI was confirmed by genomic PCR followed by Sanger sequencing. Targeted ES cells were injected into blastocysts from ICR mice, and the injected blastocysts were transferred to the uterus of pseudopregnant ICR mothers to have chimeric offspring. Obtained chimeric mice were then crossed with C57BL/6J mice, and offspring carrying the R751W allele were identified by genomic PCR followed by Sanger sequencing. Primers used for genomic PCR were as follow; Evi1_F: 5’ aacaaattcaacaagatcgcgtaaggcagc 3’, Evi1_R: 5’ gaagaaaagacattcctagtttggaccctagcag 3’. All animal experiments were approved by the Animal Care Committee of the Institute of Medical Science at the University of Tokyo (PA21-67), and were conducted following the Regulation on Animal Experimentation at The University of Tokyo based on International Guiding Principles for Biomedical Research Involving Animals.

### Cell Culture

HEK293T and Plat-E cells were cultured in Dulbecco’s Modified Eagle Medium (DMEM) (Fujifilm Wako Pure Chemical Co., Ltd., 044-29765), 10% fetal bovine serum (FBS) (Biosera), and 1% penicillin-streptomycin.

### Western blotting and Immunoprecipitation

HEK293T cells were transfected with pMYs-IRES-GFP vector, Flag-tagged wild-type or mutant MECOM, and HA-tagged wild-type or mutant CtBP1 using polyethylenimine (PEI). Cells were harvested 48 hours after transfection and lysed in Cell Lysis Buffer (Cell Signaling Technology, Danvers, MA, USA; #9803). For immunoprecipitation, cell lysates were incubated with anti-FLAG antibody (SIGMA, F3165) for 30 minutes at 4°C. Dynabeads-ProteinG (Invitrogen, USA; #10004D) was then added to the samples and incubated again for 30 minutes at 4 °C. After immunoprecipitation, samples were washed three times with Cell Lysis Buffer (Cell Signaling Technology, Danvers, MA, USA; #9803) containing 1 mM phenylmethanesulfonyl fluoride. Samples were then subjected to SDS-PAGE and transferred to a polyvinylidene fluoride membrane (Bio-Rad). The blot was incubated with anti-FLAG antibody (SIGMA, F3165) or anti-HA High affinity antibody (Roche, 12CA5) Signals were detected with ECL Western Blotting Substrate (Promega, Madison, WI, USA) and visualized with Amersham Imager 600 (GE Healthcare) or LAS-4000 Luminescent Image Analyzer (FUJIFILM).

### Immunostaining

293T cells were transfected with the pMYs-IRES-GFP vector, FLAG-tagged wild-type or mutant MECOM. Forty-eight hours after transfection, the cells were fixed with 4% paraformaldehyde for 15 min at room temperature. The cells were then permeabilized with 0.2% Triton X-100 for 5 min and blocked with BSA for 1 hour. Cells were fluorescently labeled with an anti-FLAG antibody (Sigma, F3165 or F7425) as the primary antibody and an anti-mouse antibody conjugated to Alexa Fluor 568 (Thermo Fisher, A11030) as the secondary antibody. Cell nuclei were stained with DAPI (BioLegend, catalog 422801). Fluorescence images were analyzed on an EVOS imaging system (Invitrogen).

### Luciferase assay

293T cells were seeded in 12-well culture plates at a density of 1×10^5^ cells per well 18 hours before transfection. For the luciferase assay with the proximal and distal *GATA2* promoter sequences, the cells were transfected with pGL4.1 (co-expressing Firefly Luciferase [FLuc]) containing proximal or distal promoter of *GATA2* and pGL4.74 vector (co-expressing Renilla Luciferase [RLuc]) together with pMYs-IG vector, pMYs-IG-MECOM-WT, pMYs-IG-MECOM-R750W, or pMYs-IG-MECOM-C766G using polyethylenimine (PEI). For the luciferase assay with the GATA consensus sites, the cells were transfected with pGL3 containing three repeats of the GATA consensus site (Addgene #85695) and pGL4.74 vector together with pMYS-IN vector or pMYs-NGFR-GATA2 and pMYs-IG vector, pMYs-IG-MECOM-WT, pMYs-IG-MECOM-DL/AS, pMYs-IG-MECOM-R750W, or pMYs-IG-MECOM-C766G using PEI. Cells were harvested 48 hours after transfection and were assayed for the luciferase activity using the luciferase assay system (Promega) and a luminometer (BMG LABTECH, FLUOstar OPTIMA). Promoter activity was calculated as the ratio of RLuc to FLuc.

### Viral transduction

Retroviruses for mouse cells were generated by transfecting Plat-E packaging cells with the retroviral constructs using the calcium phosphate method [44]. Retroviruses for human cells were generated by transfecting HEK293T cells with RD114 (envelope) and M57 (gag-pol) along with the retrovirus constructs using the calcium phosphate method. Lentiviruses were generated by transfecting HEK293T cells with VSVG (envelope) (Addgene, #12259) and psPAX2 (gag-pol) (Addgene, #12260) along with the lentiviral constructs using the calcium phosphate method [45]. The medium was changed after 24 hours, and the retroviral or lentiviral fluid was collected 48 hours after transfection. The retroviruses were attached to a dish coated with RetroNectin (TaKaRa, #T100B) to infect the cells.

### Flow Cytometry

Cells were stained with fluorochrome-conjugated antibodies for 30 minutes at 4 °C, washed with PBS and resuspended in PBS containing 2% FBS. Cells were then analyzed by FACS Verse (BD Bioscience). The antibodies used are provided in Supplementary Table 2.

### Colony replating, transplantation, and mast cell differentiation assays

Mouse bone marrow cells were collected from C57BL/6 (Ly5.2) female mice, 8 to 12 weeks old. Bone marrow progenitors (c-Kit^+^ cells) were selected using the CD117 MicroBead Kit (Miltenyi Biotec) and transduced with the pMYs-IRES-GFP vector, wild-type MECOM or MECOM mutants. For the colony replating assay, cells were grown in MethoCult™ M3234 (STEMCELL Technologies) with 10 ng/mL mouse stem cell factor (SCF) (R&D Systems, 455-MC), 10 ng/mL mouse granulocyte macrophage colony-stimulating factor (R&D Systems, GM-CSF) (415-ML), 10 ng/mL mouse interleukin-3 (IL-3) (R&D Systems, 403-ML) and 10 ng/mL mouse interleukin-6 (IL-6) (R&D Systems, 406-ML). For each round of plating, 1 × 10^4^ cells were plated. Colonies were counted and replated every 4 days. A colony was defined as a cluster of at least 50 cells.

For the transplantation assay, bone marrow c-Kit^+^ cells transduced with MECOM-WT, MECOM-DL/AS or MECOM-R750W were transplanted into sublethally (525 cGy) irradiated 12-week-old female C57BL/6 mice. Each mouse received 5×10^5^ cells.

For the mast cell differentiation assay, bone marrow c-Kit^+^ cells were transduced with pMYs-IG vector, MECOM-WT, MECOM-DL/AS, MECOM-R750W and MECOM-C766G. After transduction, the cells were cultured in Iscove’s Modified Dulbecco’s Medium (IMDM) (Fujifilm Wako Pure Chemical Coorperation), 10% FBS (Biosera), 1% penicillin-streptomycin supplemented with murine 1 ng/ml IL-3, 10 ng/ml IL-6 and 100 ng/ml SCF (R&D Systems). The frequency of FceR1a^+^c-Kit^+^ mast cells in the culture was evaluated by FACS every 3 days.

### Morphological evaluation

1×10^4^ cells were suspended in 100 µl PBS, and cell samples were prepared by cytospin (650 rpm, 5 min). After centrifugation, adherent cells were dried on glass slides, fixed with methanol and stained with Maygimsa stain. Hemacolor® Rapid staining of blood smears (Millipore, #111956, #111957) was used for staining. Samples were observed using an Olympus BX51TF microscope.

### Isolation and culture of human cord blood CD34^+^ cells

Human umbilical cord blood (CB) was obtained from the Kanto-Koshinetsu Umbilical Cord Blood Bank of the Japanese Red Cross Society. Mononuclear cells (MNCs) were isolated using Lymphoprep (Alere Technologies AS, Oslo, Norway). The CD34^+^ cell fraction was then isolated from the MNCs using the MidiMACS system (CD34+ Microbead Kit; Miltenyi Biotec; Bergisch Gladbach, Germany) according to the manufacturer’s protocols. CB CD34^+^ cells were incubated in StemSpanTM SFEMII (STEMCELL Technologies) supplemented with 10 ng/ml mouse Flt-3, 10 ng/ml human thrombopoietin (TPO), 10 ng/ml human stem cell factor (SCF), 10 ng/ml human interleukin-3 (IL-3) and 10 ng/ml human interleukin-6 (IL-6) (R&D Systems).

### Gata2 depletion using CRISPR/Cas9 in mouse bone marrow cells

To generate non-targeting (NT) or GATA2-targeting (sgGata2-A, B) short guide RNA (sgRNA) constructs, annealed oligos were inserted into pLentiguide-puro vector, which was obtained from Addgene (#52963). Mouse bone marrow c-Kit^+^ cells from Rosa26-LSL-Cas9 knockin mice were transduced with the sgRNAs using the lentivirus and were selected for stable expression of the sgRNAs using puromycin (1 μg/ml) in MethoCult™ M3234 (STEMCELL Technologies) supplemented with 10 ng/mL mouse SCF, 10 ng/mL mouse GM-CSF, 10 ng/mL mouse IL-3 and 10 ng/mL mouse IL-6 (R&D Systems). The efficiency of *Gata2* gene editing was assessed by ICE CRISPR Analysis Tool (https://ice.synthego.com/). Sequences for the nontargeting (NT) control and sgRNAs targeting *Gata2* are provided as follows: NT: 5’ cgcttccgcggcccgttcaa 3’, sgGata2-A: 5’ ggcgttccggcgccataagg 3’, sgGata2-B: 5’ caaccaccaccttatggcgc 3’.

### RNA-Seq

CD34^+^ human umbilical cord blood was transduced with pMYS-IRE-GFP vector, wild-type MECOM, or mutant MECOM (MECOM-R750 or MECOM-DL/AS). 48 hours after transduction, GFP^+^ cells were sorted by FACS Aria III (BD Biosciences, San Jose, CA, USA) and were cultured for 4 days as described above. Total RNA was extracted and purified from the GFP^+^ cells using the FastGene™ RNA Purification Kit (FastGene, FG-8005). Sequencing was then performed using Novaseq 6000. The fastq file was uploaded to Galaxy (https://usegalaxy.org) for analysis and mapped to the human genome (hg38) using HISAT2 [46]. Reads were then counted using Feature Count and gene expression variations were analyzed using EdgeR [47]. The data generated by Feature Count were normalized by the TPM method, and then enrichment analysis (https://maayanlab.cloud/Enrichr/) was performed. The genes whose TPM value was less than 1 in all samples were removed as low expression genes from the enrichment analysis and gene expression variation analysis.

### Quantitative RT-PCR

Mouse bone marrow cells were transduced with pMYs-IG, MECOM-WT or MECOM-R750W. After transduction, the cells were cultured in MethoCult™ M3234 (STEMCELL Technologies) with 10 ng/mL mouse SCF (455-MC), 10 ng/mL mouse GM-CSF (415-ML), 10 ng/mL mouse IL-3 (403-ML) and 10 ng/mL mouse L-6 (406-ML) (R&D Systems). Total RNA was extracted using the RNeasy Mini kit (QIAGEN) 8 days after transduction and reverse transcribed using High-Capacity cDNA Reverse Transcription Kit (Applied Biosystems). Complementary DNA (cDNA) was then subjected to quantitative RT-PCR using a SYBR Select Master Mix (Applied Biosystems). Sequences of the primers used are provided in Supplementary Table 1.

### ChIP-Seq and ChIP-qPCR

293T cells were transfected with pMYS-IG vector, MECOM-WT or MECOM-R750W with AM tag. Chromatin was then harvested, immunoprecipitated, and DNA purified using the SimpleChIP® Enzymatic Chromatin IP Kit (Magnetic Beads) (Cell Signaling, #9003). Anti-AM tag (active motif, catalog #91112) was used as the antibody for immunoprecipitation. Sequencing was performed on a next-generation sequencer from Chemical Dojin Co. The fastq file was uploaded to Galaxy (https://usegalaxy.org) for analysis and mapped to the human genome (hg38) using Bowtie2 (Galaxy Version 2.4.2+galaxy0) [48]. MACS2 (Galaxy Version 2.1.1.20160309.6) was used for peak calling [49]. ChIPseeker (Galaxy Version 1.18.0+galaxy1) was used for peak annotation [50]. BamCoverage (Galaxy Version 3.3.2.0.0), computeMatrix (Galaxy Version 3.5.1.0.0) and plotHeatmap (Galaxy Version 3.5.1.0.1) were used for sequence depth visualization [51]. The IGV tool (Version 2.8.12) was used to visualize sequence reads [52]. Homer (Version 4.10) was used to identify de novo motifs [53]. The size of the region used for motif discovery is 100 bp.

For ChIP-qPCR, purified DNA was subjected to quantitative RT-PCR after ChIP using a SYBR Select Master Mix (Applied Biosystems). The sequences of the primers used are provided in Supplemental Table 1.

## RESULTS

### Biochemical properties of wild-type and mutant MECOM

We first generated wild-type and mutant MECOM to evaluate the effect of mutations in the C-terminal ZFD and CtBP-binding sites on the MECOM function. The C-terminal ZFD MECOM mutants have the R750W (c.2248C>T) or C766G (c.2296T>G) mutation, which have been reported in MECOM-associated syndromes [35,36,37,39,41]. We also generated MECOM mutants carrying 1 (DL/AS) or 2 (AS/AS) mutations in the CtBP-binding sites. (**Figure 1A**). None of these mutations affected the expression and nuclear localization of wild-type MECOM (**Figure 1B, C**). We next examined the ability of each MECOM construct to dimerize or bind to other proteins. As previously reported [19,20,54], wild-type MECOM efficiently interacted with itself and CtBP1, whereas the DL/AS and AS/AS mutants lost the binding ability to CtBP1 (**Figure 1D**). Interestingly, the R750W mutant, but not the G766G mutant, showed the attenuated binding to CtBP1 and wild-type MECOM (**Figure 1D-E**). We also found that MECOM interacted with a hematopoietic transcription factor GATA2, which was also attenuated by the R750W mutation (**Figure 1F**). Thus, among the C-terminal ZFD mutations, only the R750W mutation reduces the protein-protein interaction of MECOM.

**Figure 1.**
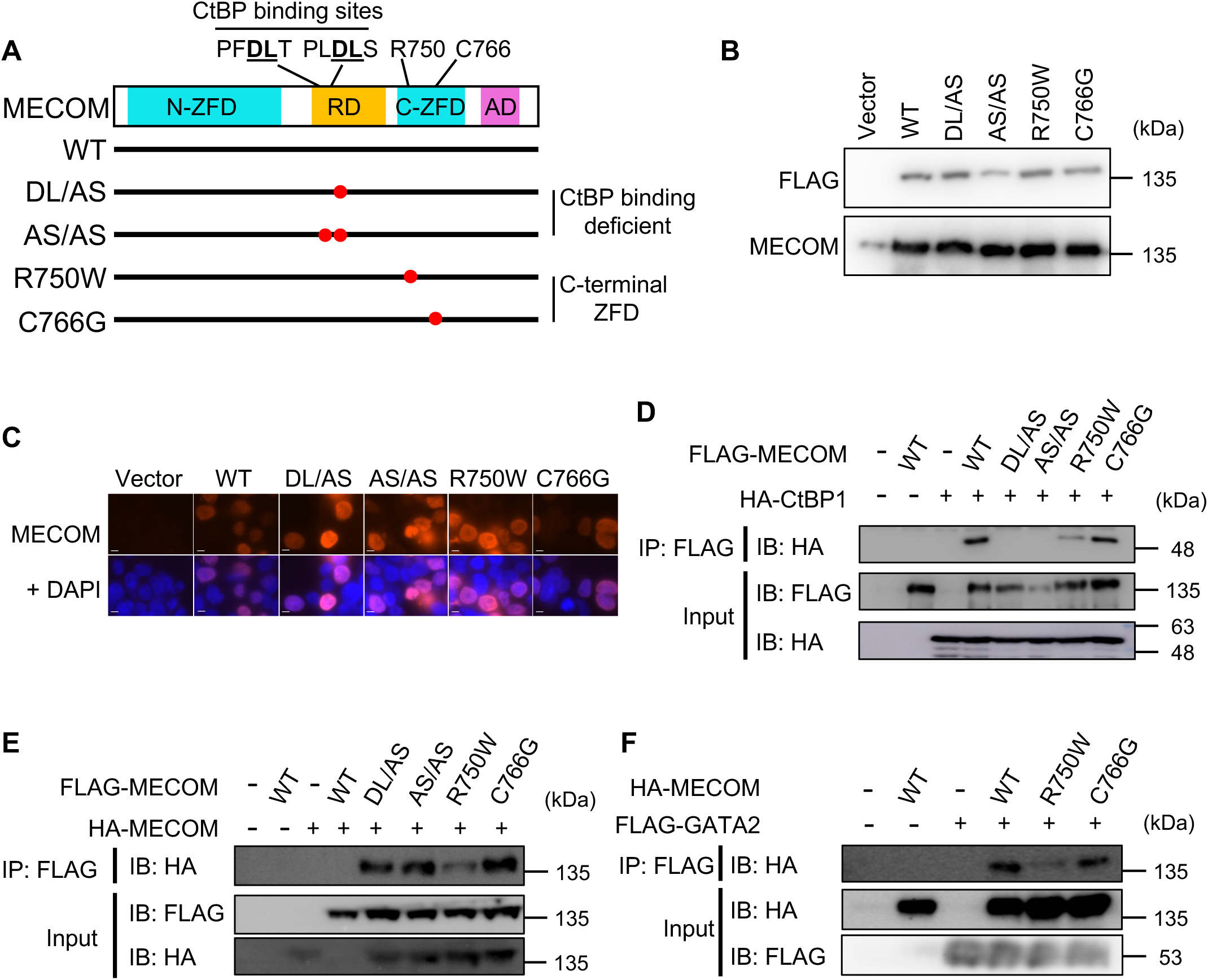
Biochemical characterization of wild-type and mutant MECOM. **A**. Schematic of wild-type (WT) and mutant MECOM. DL/AS and AS/AS lose the interaction with CtBP by carrying 1 or 2 mutations in the CtBP binding sites. R750W and C766G have mutations in the C-terminal zinc finger domain (C-ZFD), which have been reported in MECOM-associated syndromes. N-ZFD: N-terminal ZFD, RD: Repression Domain, AD: Activation Domain. **B**, **C**. 293T cells were transfected with vector or FLAG-tagged WT or mutant MECOM. MECOM levels were assessed by Western blotting with anti-FLAG or anti-MECOM antibodies (**B**). Cellular localization of WT or mutant MECOM was assessed by immunofluorescence with anti-FLAG antibody (**C**). Scale bars, 10 μm; ×1000 magnification. **D**, **E**. 293T cells were transfected with FLAG-tagged WT or mutant MECOM together with HA-tagged CtBP1 (**D**) or HA-tagged MECOM (**E**). Cell lysates were immunoprecipitated with anti-FLAG antibody, followed by the immunoblotting with anti-HA antibody. **F**. 293T cells were transfected with HA-tagged WT or mutant MECOM together with FLAG-tagged GATA2. Cell lysates were immunoprecipitated with anti-FLAG antibody, followed by the immunoblotting with anti-HA antibody.

### C-terminal ZFD and CtBP-binding site mutations abolish the MECOM function

Next, we investigated the effect of the C-terminal ZFD and CtBP-binding site mutations on the biological functions of MECOM. First, we transduced wild-type and mutant MECOM into mouse c-Kit^+^ bone marrow progenitor cells and performed a colony replating assay. Expression of each MECOM construct in mouse bone marrow cells was confirmed by western blotting (**Figure 2A)**. Consistent with previous reports, wild-type MECOM immortalized bone marrow cells and produced many immature colonies even in the third and fourth rounds of plating when the control cells stopped forming colonies. Importantly, both the C-terminal and the CtBP-binding site mutations abolished the leukemogenic activity of MECOM to promote serial replating of bone marrow cells (**Figure 2B, C**).

**Figure 2.**
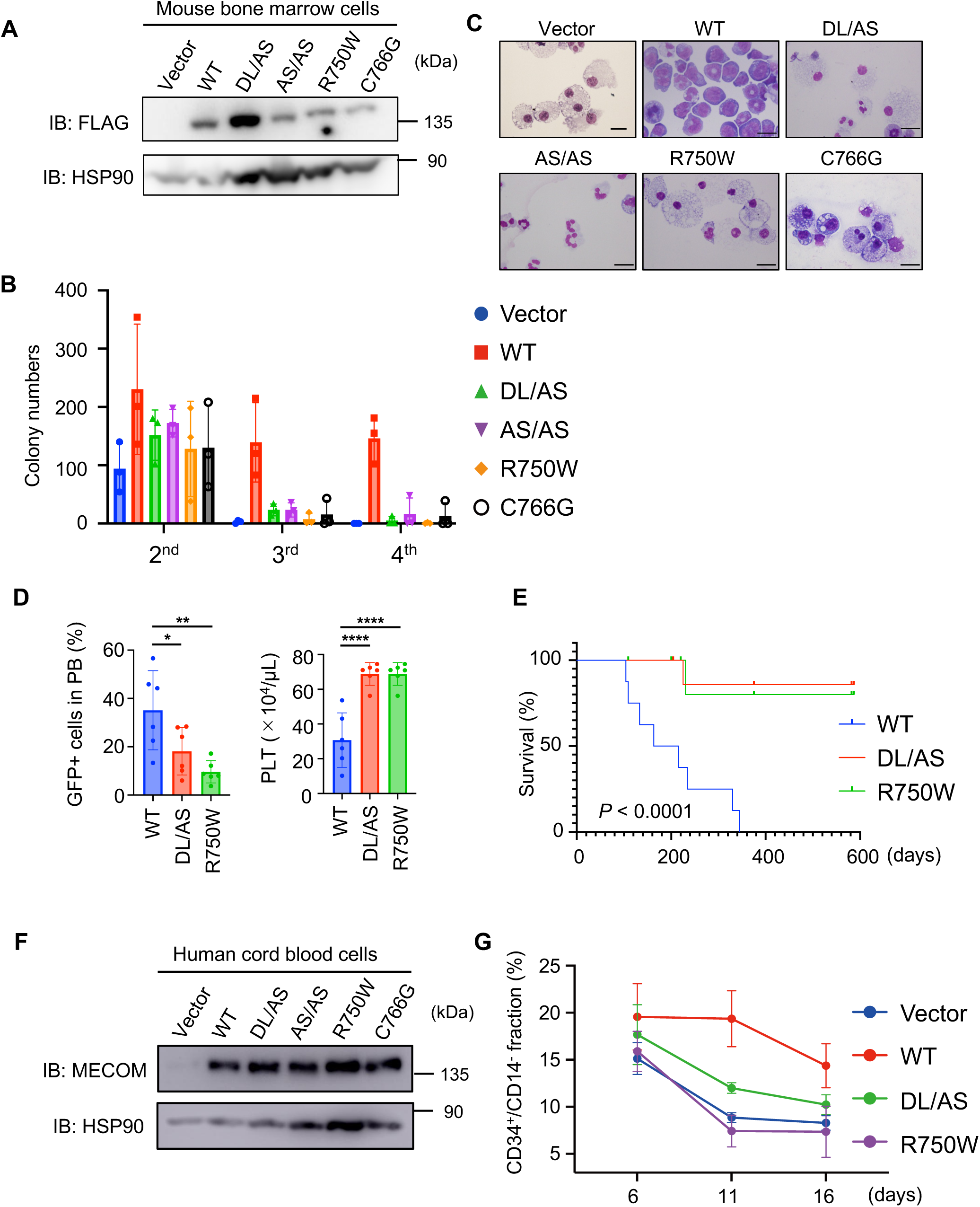
Functional analysis of wild-type and mutant MECOM. **A-C**. Mouse bone marrow c-Kit^+^ cells were transduced with vector or FLAG-tagged wild-type (WT) or mutant (DL/AS, AS/AS, R750W, C766G) MECOM. Expression of each construct was confirmed by Western blotting with anti-FLAG and anti-HSP90 antibodies (**A**). Cells were serially replated every four days. Colonies were counted before replating (**B**, data are shown as mean with SD, *P < 0.05, **P < 0.01, two-way ANOVA with Dunnett‘s multiple comparisons test, n = 3). Representative images of each colony at the fourth round are also shown (**C**, Scale bars, 20 μm; ×1000 magnification). **D**, **E**. Mouse bone marrow c-Kit^+^ cells were transduced with WT or mutant (DL/AS or R750W) MECOM coexpressing GFP and transplanted into recipient mice. Peripheral blood data 1 month after transplantation (**D,** data are shown as mean with SD, *P < 0.05, **P < 0.01, one-way ANOVA with Dunnett‘s multiple comparisons test, n = 6) and survival curves (**E**, n = 8 each, log-rank test) are shown. **F**, **G**. Human cord blood (CB) CD34^+^ cells were transduced with vector, WT, DL/AS, AS/AS, R750W and C766G (coexpressing GFP). Expression of each construct was confirmed by Western blotting with anti-MECOM and anti-HSP90 antibodies (**F**). Cells were cultured in myeloid skewing medium and analyzed by FACS to measure the frequency of CD34^+^/CD14^-^ cells (**G**, data are shown as mean with SD, **P < 0.01, ****P < 0.0001, two-way ANOVA with Dunnett‘s multiple comparisons test).

We then transplanted mouse bone marrow c-kit^+^ cells transduced with wild-type MECOM, MECOM-DL/AS, or MECOM-R750W into sublethally irradiated recipient mice. The mice transplanted with wild-type MECOM-expressing cells had more GFP^+^ leukemic cells, showed a significant reduction in platelet count one month after transplantation (**Figure 2D**), and developed AML within one year (**Figure 2E**, **Supplementary Figure 1A-C**). In contrast, neither MECOM-R750W nor MECOM-DL/AS induced AML development, suggesting that both types of MECOM mutants lose the leukemogenic activity *in vivo* (**Figure 2E**).

We also evaluated the effect of wild-type and mutant MECOM on the self-renewal capacity of human HSCs. Cord blood (CB) CD34^+^ cells were transduced with vector, wild-type MECOM, MECOM-DL/AS or MECOM-R750W, and were cultured *in vitro* to monitor the frequency of CD34^+^/CD14^-^ cells chronologically by FACS analysis. Expression of each MECOM construct in CB CD34^+^ cells was confirmed by Western blotting (**Figure 2F**). Wild-type MECOM inhibited the myeloid maturation of CB cells, as evidenced by the increase of CD34^+^/CD14^-^ immature cells in culture (**Figure 2G**). In contrast, neither MECOM-DL/AS nor MECOM-R750W inhibited the myeloid differentiation of CB cells. MECOM-G766G also failed to inhibit myeloid maturation (**Supplementary Figure 2A-B**). Taken together, we concluded that the C-terminal ZFD mutations and the CtBP-binding site mutations are loss-of-function mutations.

### C-terminal ZFD mutations abolish DNA binding and transcriptional activity of MECOM

Since the R750W and C766G mutations are located in the C-terminal ZFD, which has DNA-binding ability, we speculated that the C-terminal ZFD mutations may inhibit the binding of MECOM to DNA. To test this hypothesis, we transfected HEK293T cells with vector, wild-type MECOM and MECOM-R750W tagged with AM-tag and performed ChIP-seq analysis (**Supplementary Figure 3A**). The AM-tag is specifically designed to provide low background signal in ChIP experiments. Wild-type MECOM bound mainly to the promoter regions of its target genes, including *GATA2* and *PBX1* (**Figure 3A-C, Supplementary Figure 3B**). Notably, the most significantly enriched motif resembles the known binding motif of the C-terminal ZFD (5’-GAAGATGAG-3’), but not that of the N-terminal ZFD (5’- GA(C/T)AAGA(T/C)AAGATAA-3’) of MECOM (**Figure 3D**). MECOM-R750W also bound to similar genomic regions as wild-type MECOM, but less efficiently (**Figure 3B**, **C, Supplementary Figure 3B**). The reduced DNA binding ability of MECOM-R750W and MECOM-C766G on the *GATA2* promoters was also confirmed by ChIP-qPCR analysis (**Figure 3E, Supplementary Figure 3C**). We then performed luciferase reporter assays using reporters containing the proximal and distal GATA2 promoter sequences, which have been shown to be activated by wild-type MECOM [5]. Consistent with the reduced DNA binding ability, MECOM-R750W and MECOM-C766G showed significantly reduced transcriptional activity compared to wild-type MECOM (**Figure 3F**). These data suggest that MECOM binds to DNA mainly through the C-terminal ZFD, which is inhibited by the mutations associated with the MECOM-associated syndromes.

**Figure 3.**
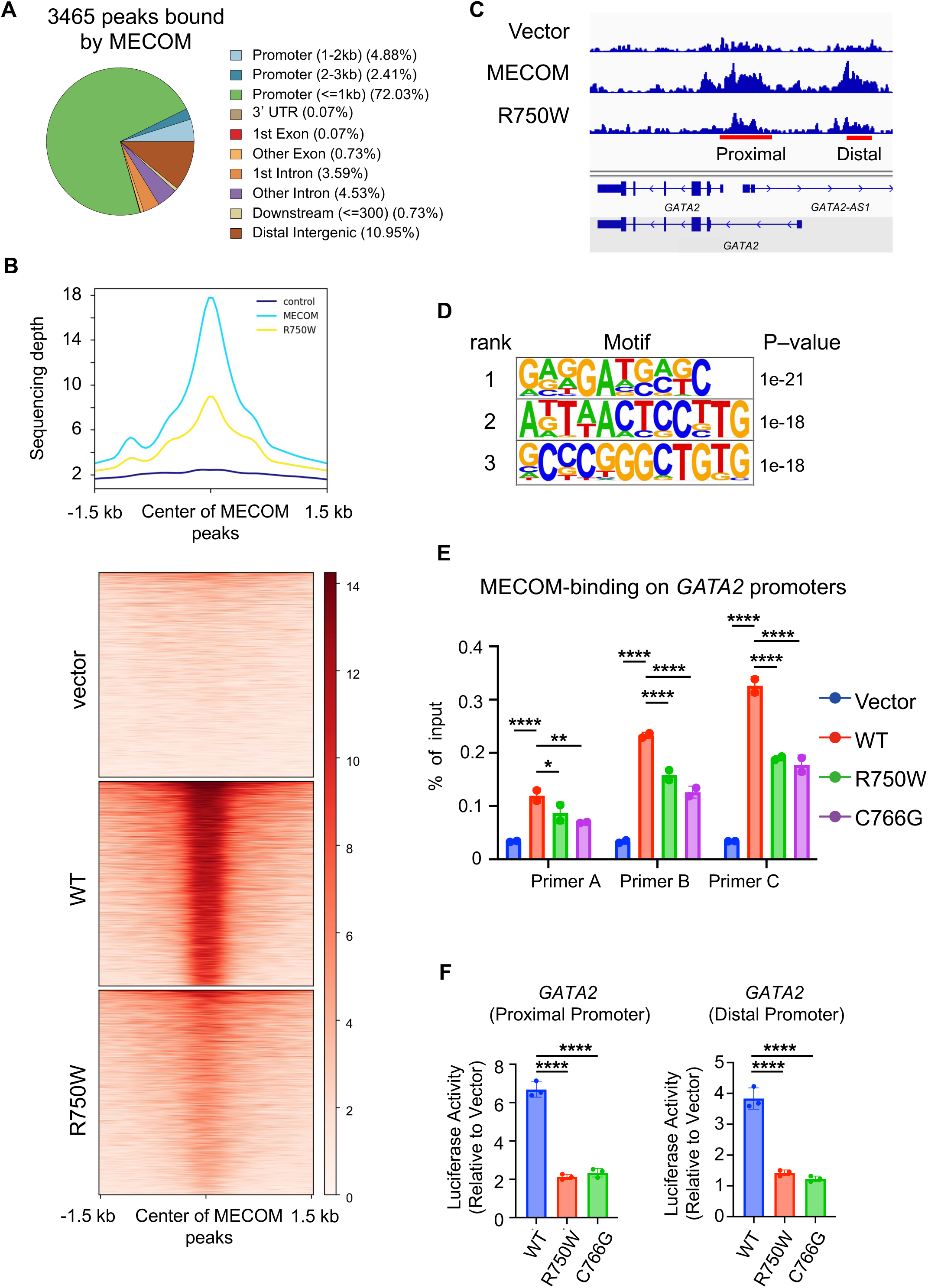
Reduced DNA binding ability of mutant MECOM with C-terminal ZFD mutations. **A-C**. 293T cells were transfected with vector, AM-tagged wild-type (WT) MECOM or MECOM-R750W, immunoprecipitated with anti-AMtag antibody, followed by sequencing analysis. Genomic distribution of anti-AMtag ChIP-seq peaks (**A**), density maps of ChIP-seq peaks (**B**), and sequence reads at the proximal and distal GATA2 promoter (**C**) are shown. **D**. De novo motif analysis of wild-type MECOM ChIP-seq peaks (a 70bp DNA sequence extracted from each significant peak) using the Homer application (ver4.10). **E**. Binding of MECOM-WT and MECOM-R750W to the proximal GATA2 promoter was assessed by ChIP-qPCR using three independent primers (see also **Supplementary Figure 3C**). Data are shown as mean with SD, *P < 0.05, **P < 0.01, ****P < 0.0001, two-way ANOVA with Dunnett‘s multiple comparisons test, n=2). **F**. 293T cells were transfected with vector, FLAG-tagged wild-type (WT) or mutant (R750W, C766G) MECOM together with the reporter plasmid containing the proximal or distal GATA2 promoter. Relative luciferase activity is expressed as the ratio of luciferase activity to vector control (mean with SD, ****P < 0.0001, one-way ANOVA with Dunnett‘s multiple comparisons test, n=3).

### Wild-type MECOM, but not the mutant MECOM, represses expression of mast cell-associated genes

Next, we performed RNA-seq using human CB CD34^+^ cells transduced with vector, wild-type MECOM, MECOM-DL/AS or MECOM-R750W (coexpressing GFP). Hierarchical clustering and principal component analysis revealed that the CB cells expressing each MECOM construct had unique expression profiles (**Figure 4A, Supplementary Figure 4A**). As MECOM-repressed genes, we identified 120 and 255 genes that were downregulated by wild-type MECOM but not by MECOM-R750W or MECOM-DL/AS, respectively (**Figure 4B, Supplementary Table 3**). We also identified 194 and 447 genes that were upregulated by wild-type MECOM but not by MECOM-R750W or MECOM-DL/AS, respectively, as MECOM-activated genes (**Supplementary Figure 4B, C Supplementary Table3**). Interestingly, enrichment analysis of the MECOM-repressed genes revealed the significant downregulation of mast cell-associated genes in wild-type MECOM-expressing cells (**Figure 4C**). We confirmed the downregulation of several mast cell genes (HDC, CPA3, TPSB2 and MITF) in wild-type MECOM-expressing CB cells, which was partially restored in mutant MECOM-expressing cells (**Figure 4D**).

**Figure 4.**
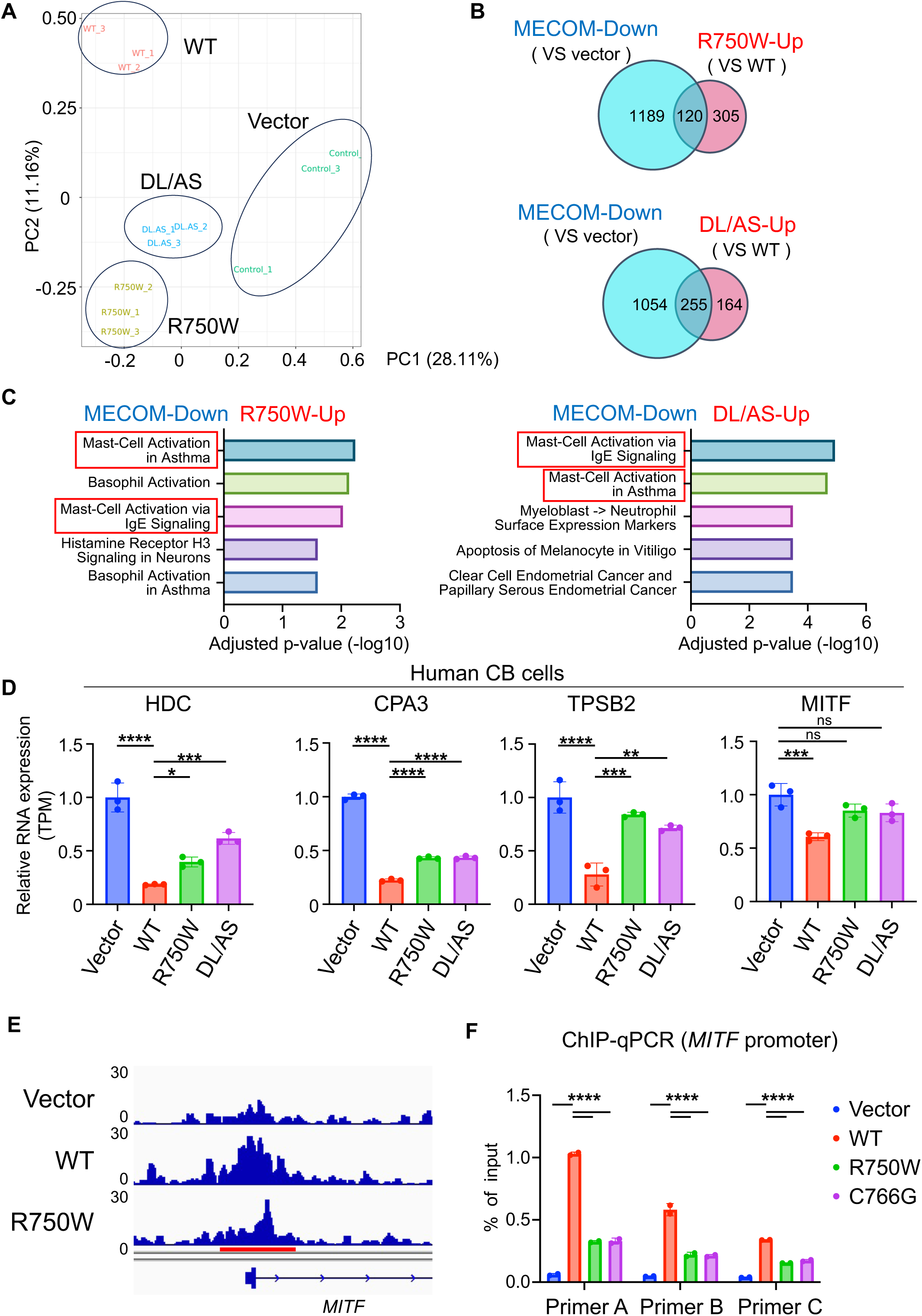
Mast cell genes are repressed by wild-type, but not mutant, MECOM. **A-D**. Human cord blood (CB) CD34^+^ cells were transduced with vector, wild-type (WT) or mutant (DL/AS, R750W) MECOM coexpressing GFP. After 4 days of culture in stem cell medium, RNA was extracted from GFP^+^ cells and subjected to RNA-seq. (**A**) Principal Component Analysis (PCA) of RNA-seq data. (**B**) Venn diagram showing the genes downregulated in MECOM-WT-transduced CB cells compared to vector-transduced CB cells (blue circle) and genes upregulated in MECOM-R750W- (upper panel) or MECOM-DL/AS- (lower panel) transduced CB cells compared to MECOM-WT-transduced CB cells (red circle). Genes with FDR < 0.05 were considered as up- or down-regulated genes. (**C**) The 120 (for R750W) and 255 (for DL/AS) overlapping genes were used for enrichment analysis using Elsevier Pathway Collection. The mast cell-related gene sets are highlighted by the red boxes. (**D**) Transcripts Per Million (TPM) of mast cell-related genes (HDC, CPA3, TPSB2 and MITF) in vector, WT, or mutant MECOM-transduced CB cells. *P < 0.05, **P < 0.01, ***P < 0.001, ****P < 0.0001, one-way ANOVA with Dunnett’s multiple comparisons test. n=3. **E**. 293T cells were transfected with vector, AM-tagged wild-type (WT) MECOM or MECOM-R750W, immunoprecipitated with anti-AMtag antibody, followed by sequencing analysis. Sequence reads at the MITF promoter are shown. **F**. 293T cells were transduced with vector, AM-tagged MECOM-WT, R750W or C766G, immunoprecipitated with anti-AMtag antibody, followed by qPCR using three primers designed for different regions in the MITF promoter (see also **Supplementary Figure 5B**). Data are shown as mean with SD, ****P < 0.0001, two-way ANOVA with Dunnett‘s multiple comparisons test, n = 2).

We then combined ChIP-seq and RNA-seq data to identify direct transcriptional targets of MECOM in human myeloid progenitor cells. Among 2286 MECOM-bound genes, we identified 15 and 21 genes whose expression was up- or down-regulated, respectively, only by wild-type MECOM (**Supplementary Figure 5**). One gene of interest was MITF, a basic helix-loop-helix leucine zipper transcription factor that plays a critical role in mast cell development (**Figure 4D, Supplementary Figure 5A**). We confirmed the strong or weak binding of wild-type and mutant MECOM to the promoter region of MITF by ChIP-Seq and ChIP-qPCR (**Figure 4E, F, Supplementary Figure 5B**). We also found that forced expression of wild-type MECOM, but not MECOM-R750W, induced Mitf downregulation in mouse bone marrow c-kit^+^ cells (**Supplementary Figure 5C**). Thus, MITF, a key regulator of mast cell differentiation, is a direct transcriptional target of MECOM. Collectively, these results suggest that wild-type MECOM suppresses the expression of mast cell-associated genes whose activity is reduced by mutations in the C-terminal ZFD or CtBP binding sites.

### MECOM suppresses mast cell differentiation and promotes leukemogenesis by antagonizing GATA2

Next, we investigated the potential role of MECOM in the mast cell differentiation. Mouse bone marrow c-Kit^+^ cells were transduced with vector, wild-type and mutant (DL/AS, R750W, C766G) MECOM, and were cultured *in vitro* with cytokines to induce mast cell differentiation. Most of the vector-transduced cells became Fcer1a^+^c-Kit^+^ mast cells at day 12. In contrast, the frequency of FceR1a^+^c-Kit^+^ mast cells in the wild-type MECOM-expressing cells was less than 10 %, indicating that MECOM is a suppressor of mast cell differentiation. Importantly, all the MECOM mutations in the C-terminal ZFD or CtBP-binding sites abolished this suppressive activity of MECOM on mast cell maturation (**Figure 5A, B**). We also assessed the effect of forced expression of wild-type and mutant MECOM on GATA2-induced mast cell maturation. Mouse bone marrow c-Kit^+^ cells were transduced with the MECOM constructs together with GATA2 and grown in the mast cell cultures. As expected, GATA2 promoted mast cell differentiation, which was efficiently suppressed by wild-type MECOM. On the other hand, all MECOM mutants (MECOM-R750W, C766G and DL/AS) suppressed GATA2-induced mast cell maturation only partially, again confirming that they are the loss-of-function mutations (**Figure 5C, D**).

**Figure 5.**
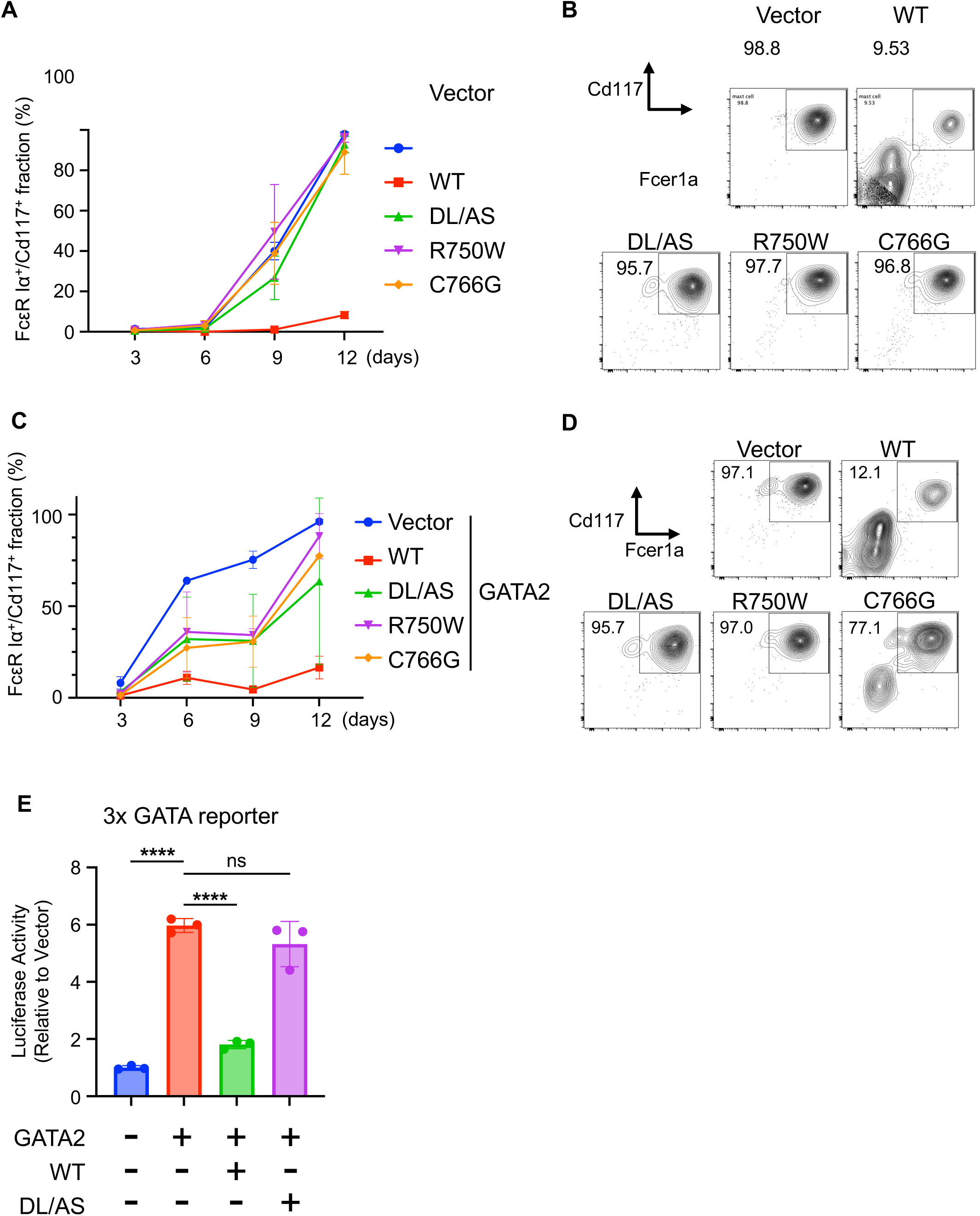
MECOM inhibits GATA2 activity and mast cell differentiation. **A, B**. Mouse bone marrow c-Kit^+^ cells were transduced with vector, wild-type (WT) or mutant (DL/AS, R750W, C766G) MECOM (coexpressing GFP) and cultured in the mast cell-inducing culture. The frequency of the mast cell fraction (c-Kit^+^/Fcer1a^+^) in GFP^+^ cells was measured chronologically by FACS (**A**). n=2 for each group. Representative FACS plots at day 12 are also shown (**B**). Numbers indicate the frequency (%) of c-Kit^+^/Fcer1a^+^ fraction in GFP^+^ cells. **C**, **D**. Mouse bone marrow c-Kit^+^ cells were transduced with vector, WT or mutant (DL/AS, R750W, C766G) MECOM (coexpressing GFP) together with vector or GATA2 (coexpressing NGFR) and cultured in the mast cell inducing culture. The frequency of the mast cell fraction (c-Kit^+^/Fcer1a^+^) in GFP^+^NGFR^+^ cells was measured chronologically by FACS (**C**). n=2 for each group. Representative FACS plots at day12 are also shown (**D**). Numbers indicate the frequency (%) of c-Kit^+^/Fcer1a^+^ fraction in GFP^+^ cells. **E**. 293T cells were transfected with vector, WT or mutant (DL/AS, R750W, C766G) MECOM together with GATA2 and the reporter plasmid containing 3x GATA sequences. Relative luciferase activity is shown as the ratio of luciferase activity to vector control (mean with SD, ****P < 0.0001, one-way ANOVA with Dunnett‘s multiple comparisons test, n=3). ns: not significant.

Since GATA2 is a transcription factor that determines mast cell identity, we then examined a potential inhibitory effect of MECOM on GATA2-mediated transcription using a luciferase reporter containing 3x GATA consensus sequences. MECOM efficiently suppressed the GATA2-induced activation of the GATA2-reporter, whereas MECOM-DL/AS did not (**Figure 5E**). Thus, MECOM inhibits GATA2-mediated transcription through the recruitment of the corepressor CtBP, thereby suppressing mast cell differentiation.

### GATA2 depletion partially restores leukemogenic activity of mutant MECOM

We next investigated the possibility that the MECOM-mediated GATA2 suppression also contributes to MECOM-induced leukemogenesis. Mouse bone marrow c-Kit^+^ cells were collected from Cas9 knockin mice and transduced with vector, wild-type MECOM or MECOM-R750W together with single guide RNAs (sgRNA) targeting C-terminal zinc finger region of Gata2, where the mutations found in MonoMAC syndromes are concentrated [55]. These cells were serially replated in semisolid media (**Figure 6A**). At the third passage, vector-transduced cells produced only a few numbers of colonies even with the GATA2 sgRNAs, indicating that GATA2 depletion alone was not sufficient to enhance the self-renewal of bone marrow cells. In addition, GATA2 depletion showed no effect on the colony numbers of MECOM-expressing cells, indicating that wild-type MECOM does not need further reduction of GATA2 function to immortalize mouse bone marrow cells. Interestingly, GATA2 depletion restored colony forming activity of the mutant MECOM (R750W and DL/AS)-expressing cells at the third passage (**Figure 6B)**. *GATA2* gene was indeed cleaved in the colony forming cells expressing MECOM-R750W while no editing was observed in cells expressing wild-type MECOM (**Supplementary Figure 6**). These results suggests that the C-terminal ZFD mutants of MECOM lost the transforming activity partly because they are unable to inhibit GATA2 function.

**Figure 6.**
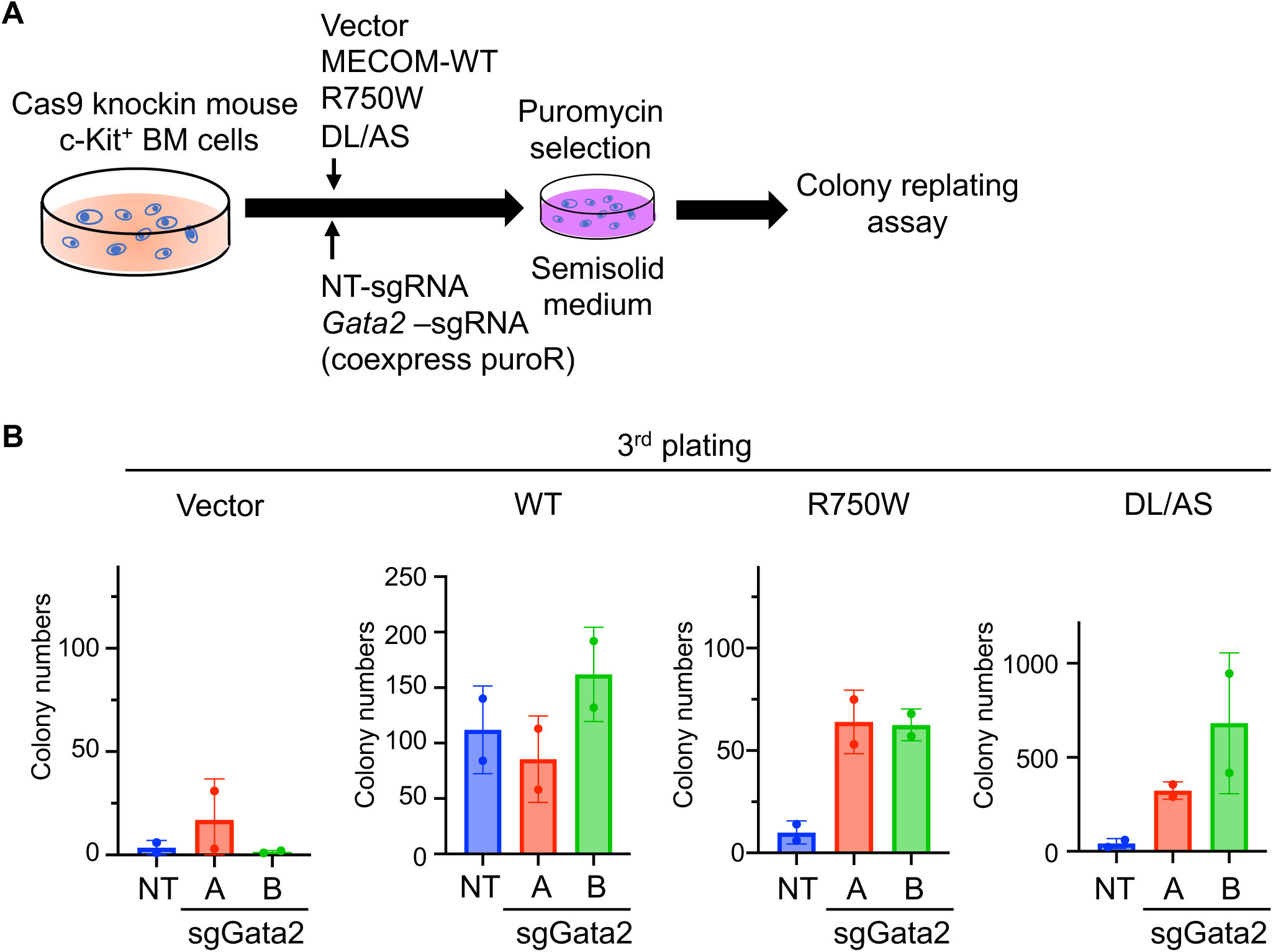
GATA2 depletion partially restores the colony replating ability of mutant MECOM. **A**. Experimental scheme used in **B**. Mouse bone marrow c-Kit^+^ cells were transduced with vector, wild-type (WT), R750W or DL/AS together with non-targeting (NT) or two independent Gata2-targeting sgRNAs [sgGata2-A and sgGata2-B coexpressing puromycin-resistant gene (PuroR)]. The sgRNA-transduced cells were selected with 1 μg/ml puromycin and were serially replated every four days. **B**. Total colony numbers at the third plating are shown.

### Hematopoietic dysregulation in *MECOM*^WT/R751W^ mice

To investigate the effect of the R750W mutation on the *in vivo* function of MECOM, we generated knockin mice harboring a point mutation (Mecom: [NP_001415378.1], R751W, [NM_001428449.1], c.C3041T), corresponding to the MECOM-R750W identified in patients with MECOM-associated syndromes. Homozygous mutant mice died before embryonic day 14.5 (E14.5) (**Figure 7A**). Heterozygous mutant mice (MECOM^WT/R751W^ mice) grew normally and showed no obvious hematopoietic defects in the peripheral blood during one and a half years of observation (**Supplementary Figure 7A**). We next analyzed the frequency of various hematopoietic fractions in fetal livers and bone marrows derived from MECOM^WT/R751W^ mice and their littermates. We found a significant decrease in the frequency of Lineage^-^Sca-1^+^c-Kit^+^ (LSK) fraction containing HSCs in the fetal liver at E14.5 (**Figure 7B**), and a decrease in the frequency of SLAM-LSK fraction containing long-term HSCs (**Figure 7C**) in the adult bone marrow of MECOM^WT/R751W^ mice. B220^+^ B-cells were also modestly reduced in the bone marrow of MECOM^WT/R751W^ mice, while the frequencies of Gr-1^+^ and Ter119^+^ cells were not changed significantly (**Figure 7D**). Thus, the MECOM^WT/R751W^ mice recapitulate the HSC defects and B-cell deficiency observed in some patients with MECOM-associated syndromes. However, the relatively mild phenotype of the MECOM^WT/R751W^ mice also suggests that other factors are required for the development of MECOM-associated syndromes.

**Figure 7.**
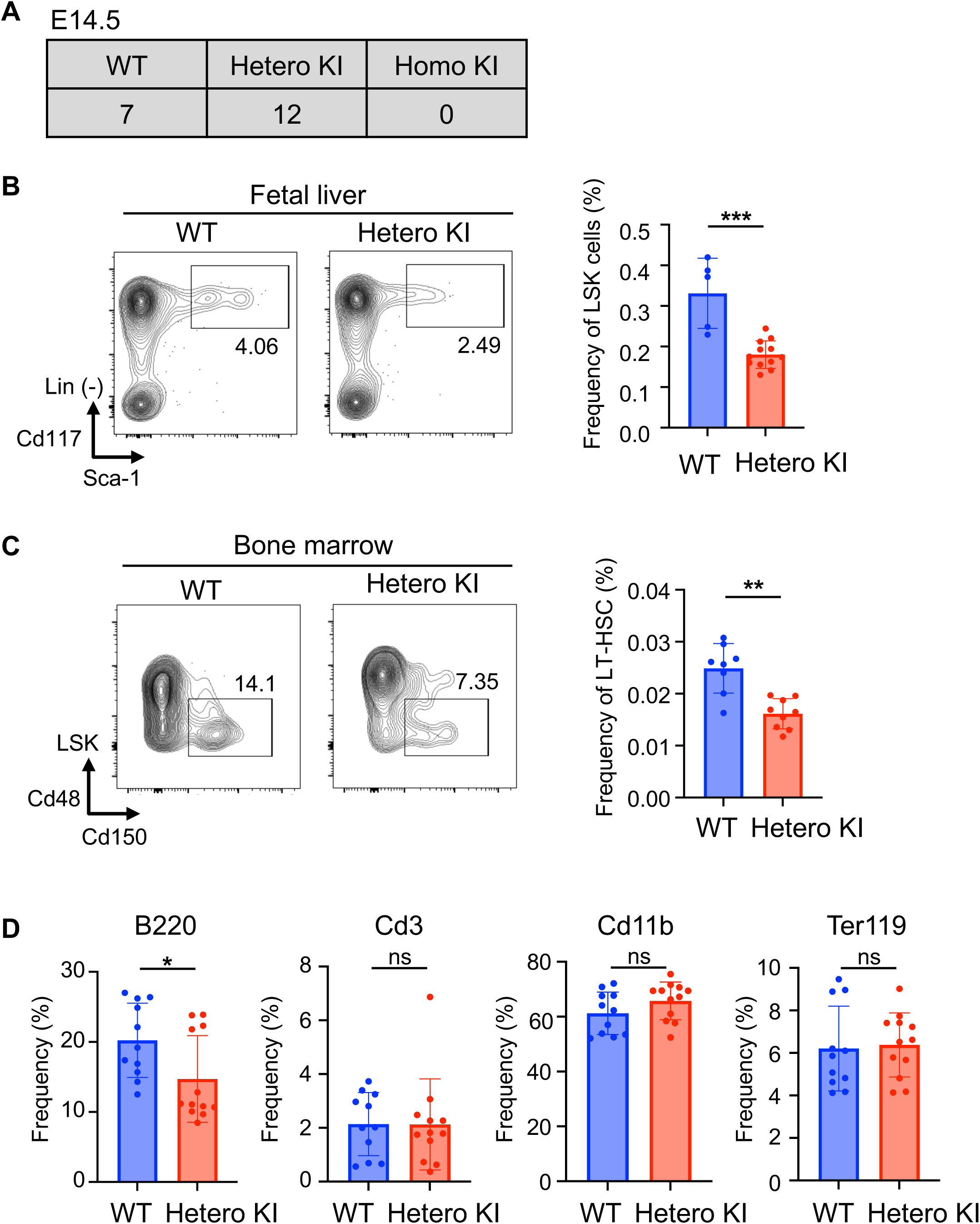
Analysis of MECOM^WT/R751W^ knockin mice. **A.** Genotypes of litters obtained by intercrossing MECOM^WT/R751W^ mice at embryonic (E) day 14. **B**. Frequency of Lineage^-^Sca1^+^c-Kit^+^ (LSK) cells in fetal livers of wild-type (WT) and MECOM^WT/R751W^ mice at E14.5. Representative FACS plots (left, numbers indicate the frequency of LSK cells) and their quantification (right). Data are presented as mean with SD, ***P < 0.001, unpaired and two-tailed Mann-Whitney test, WT mice: n = 5, MECOM^WT/R751W^ mice: n = 12). **C**. Frequency of SLAM (Cd150^+^Cd48^-^) LSK cells in bone marrow of 2-month-old wild-type and MECOM^WT/R751W^ mice. Representative FACS plots (left, numbers indicate the frequency of SALM-LSK cells) and their quantification (right). Data are presented as mean with SD, **P < 0.01, unpaired and two-tailed Mann-Whitney test, WT: n = 8, MECOM^WT/R751W^ mice: n = 9). **D**. Frequency of B220^+^, Cd3^+^, Cd11b^+^ and Ter119^+^ cells in bone marrow of mice analyzed in (**C**). Data are presented as mean with SD, *P < 0.05, unpaired and two-tailed Mann-Whitney test, WT: n = 11, MECOM^WT/R751W^ mice: n = 12)

## DISCUSSION

MECOM is a transcription factor with N- and C-terminal ZFDs that mediate DNA binding. Although recent genetic analyses have revealed that mutations are clustered at the C-terminal ZFD of *MECOM* in the MECOM-associated syndromes [35,36,37,38,39,40,41], how the C-terminal ZFD mutations affect the MECOM’s function was unknown. In this study, we showed that the two mutant MECOM with the C-terminal ZFD mutations, MECOM-R750W and C766G, have reduced DNA binding ability and transcriptional activity. The reduced binding of MECOM-R750W to almost all the MECOM-target genes strongly suggest that C-terminal ZFD is a major region to mediate the MECOM-DNA interaction in cells. The R750W mutation also reduces the protein-protein interaction of MECOM, which could contribute to the development of MECOM-associated syndromes. However, given that another MECOM mutant C766G retains normal protein-protein interaction while completely loses its transcriptional and transforming ability, the loss of DNA binding is probably a major reason for the loss-of-function of the C-terminal ZFD mutations.

Our study also revealed complex regulation of GATA2 by MECOM. Consistent with previous reports [5], MECOM activates luciferase reporters containing the proximal and distal promoters of GATA2, indicating that MECOM directly upregulates GATA2 transcription. On the other hand, MECOM inhibits GATA2-mediated activation of the reporter containing 3x GATA sequences, suggesting that MECOM competes with GATA2-induced transcription. Thus, MECOM increases GATA2 expression but at the same time inhibits GATA2-mediated transcription, thereby finetunes GATA2 activity. In myeloid progenitor cells, MECOM-induced “inhibition” of GATA2 appears to be important for suppression of myeloid differentiation and leukemogenesis. We found that the impaired leukemogenicity of MECOM-R750W mutant was partially reversed by GATA2 depletion, and that MECOM overexpression inhibits GATA2-induced mast cell maturation. Furthermore, a previous report showed that GATA2 haploinsufficiency accelerates MECOM-driven leukemogenesis [33,34]. These findings suggest that MECOM promotes leukemogenesis through “inhibition”, not “activation”, of GATA2 in myeloid progenitors. In contrast, previous reports have shown that MECOM promotes early hematopoiesis through GATA2 upregulation [5,30]. Therefore, MECOM-induced GATA2 “activation” may be important for HSC expansion during embryogenesis. The precise role of MECOM-GATA2 axis in diverse biological contexts warrants further investigation.

To confirm that C-terminal ZFD mutations are loss-of-function mutations *in vivo*, we generated a knockin mouse harboring a murine MECOM (mMECOM)-R751W [corresponding to human MECOM (hMECOM)-R750W] mutation. Consistent with the phenotypes of previously generated MECOM knockout mice [3,4,5,56], homozygous mutant mice (MECOM^R751W/R751W^) were embryonic lethal and heterozygous mutant mice (MECOM^WT/R751W^) showed a substantial reduction of HSCs in both fetal liver and adult bone marrow. The MECOM^WT/R751W^ mice also showed a modest but significant decrease in B cells, which is observed in a subset of MECOM-associated syndromes. Recently, another knockin mouse with a mMECOM-H752R (corresponding to hMECOM-H751R) mutation was generated, which also showed reduced HSCs in the bone marrow [42]. Thus, the two C-terminal ZFD mutations are indeed loss-of-function mutations and can induce the hematopoietic abnormalities in mice. The slight phenotypic difference between our hMECOM-R751W mice and the mMECOM-H752R mice (MECOM^WT/752R^ mice showed thrombocytopenia while our mice did not) suggests that each MECOM mutant has its own activity, which may contribute to the diverse representation of MECOM-associated syndromes. In addition, the relatively weak phenotypes of these two mutant MECOM knockin mice suggest that other factors, such as additional mutations, determine the phenotypes of MECOM-associated syndromes.

In summary, we have shown that the C-terminal ZFD mutations found in MECOM-associated syndromes are loss-of-function mutations with reduced DNA-binding capacity. MECOM antagonizes GATA2 to inhibit myeloid maturation and promote myeloid leukemogenesis through the C-terminal ZFD-mediated DNA binding and CtBP interaction. In addition, our mutant MECOM knockin mice will be a useful model to study MECOM-associated syndromes.

## Acknowledgements

We thank Shiori Shikata and Akiho Tsuchiya for technical assistance. We also thank the Flow Cytometry Core and the Mouse Core at The Institute of Medical Science, The University of Tokyo. This work was supported by Grant-in-Aid for Scientific Research (B) (22H03100, SG), Grant-in-Aid for Scientific Research on Innovative Areas (Research in a proposed research area) (21H00274, SG), Fostering Joint International Research (B) (22KK0127, SG), AMED under Grant Number (22ck0106644s0202 and 23ama221514h0002, SG), the 36^th^ Novartis Research Grant (SG), research grants from The Japanese Society of Hematology (SG, TK), Grant-in-Aid for Scientific Research (A) (20H03537, TK), JSPS KAKENHI Grant Number JP24K19216 (KY), a research grant from The Mochida Memorial Foundation for Medical and Pharmaceutical Research (KY) and a research grant from The Uehara Memorial Foundation.

## Authors’ contributions

K.I. designed and performed experiments, analyzed the data, and wrote the paper. M.N. designed and performed experiments and analyzed the data. J.N., S.A., T.I. and T.Y. assisted in the experiments and analyzed the data. M.O. and Y.Y. developed and provided chimera MECOM^WT/R751W^ mice. T.K. and K.Y. conceived the project and analyzed the data. S.G. conceived the project, designed experiments, analyzed the data and wrote the paper.

## Disclosure of Conflicts of Interest

All authors declare no competing financial interests with the contents of this article.

## Supplementary figure legends

**Supplementary figure 1: Flow cytometric profiles of MECOM-driven leukemia. A**, **B.** FACS analysis of GFP^+^ (MECOM-transduced) cells in bone marrow (**A**) and spleen (**B**).

**Supplementary figure 2: Effects of wild-type and mutant MECOM on self-renewal of cord blood cells.**

**A**. Human cord blood (CB) CD34^+^ cells were transduced with vector, wild-type (WT) or mutant (R750W, C766G) MECOM coexpressing GFP. The cells were cultured in myeloid skewing medium and analyzed by FACS to measure the frequency of CD34^+^/CD14^-^ cells. See also Figure 2H. **B.** Representative images of CB cells transduced with the indicated construct at day12.

**Supplementary figure 3: ChIP-seq analysis of 293T cells expressing wild-type (WT) MECOM or MECOM-R750W.**

**A.** Expression of AM-tagged MECOM-WT and MECOM-R750W was confirmed by Western blotting. **B**. Visualization of MECOM-WT and MECOM-R750W bound to the *PBX1.* **C**. Scheme of the position of the primers (Primer A, B, C) used for the ChIP-qPCR for the *GATA2* proximal promoter.

**Supplementary figure 4: RNA-seq analysis of wild-type and mutant MECOM-transduced human cord blood cells.**

**A.** Hierarchical clustering of the RNA-Seq data from human cord blood (CB) cells transduced with vector, wild-type (WT) or mutant (DL/AS, R750W) MECOM. n = 3 for each group. **B.** (Left) Venn diagram showing the genes upregulated in MECOM-WT transduced CB cells compared to vector-transduced CB cells (red circle) and genes downregulated in MECOM-R750W transduced CB cells compared to MECOM-WT transduced CB cells (blue circle). Genes with FDR<0.05 were considered as up- or down-regulated genes. (Right) The 194 overlapping genes were used for enrichment analysis using Elsevier Pathway Collection. **C.** (Left) Venn diagram showing the genes upregulated in MECOM-WT transduced CB cells compared with vector-transduced CB cells (red circle) and genes downregulated in MECOM-DL/AS transduced CB cells compared with MECOM-WT transduced CB cells (blue circle). Genes with FDR<0.05 were considered as up- or down-regulated genes. (right) The 447 overlapping genes were used for enrichment analysis using Elsevier Pathway Collection.

**Supplementary figure 5: Identification of MECOM target genes.**

**A**. Venn diagram showing the overlap between 2286 genes bound by MECOM (green circle) and 194 genes upregulated by MECOM (red circle: left) or 120 genes downregulated by MECOM (red circle: right). The 15 MECOM-activated and 21 MECOM-repressed genes are shown in the lists. **B**. Scheme of the position of the primers (Primer A, B, C) used for the ChIP-qPCR for the MITF promoter. **C**. Mouse bone marrow c-Kit^+^ cells were transduced with vector, MECOM-WT or MECOM-R750W and were cultured in semi-solid medium. RNA was extracted on day 8 and Mitf levels were analyzed by qRT-PCR. Results were normalized to Actb, with the relative mRNA level in vector-transduced cells set to 1. Data are presented as mean with SD of two biologically independent experiments. **P < 0.01, one-way ANOVA with Dunnett’s multiple comparisons test (n=2).

**Supplementary figure 6: Confirmation of Gata2 depletion in MECOM-R750W transduced cells**

**A**, **B.** DNA was extracted from Cas9^+^ mouse bone marrow cells transduced with wild-type MECOM (**A**) or MECOM-R750W (**B**) together with non-targeting (NT) or two independent Gata2-targeting sgRNAs (sgGata2-A or sgGata2-B). Sequences near the PAM sequence (red line) were evaluated. Note that R750W/sgGata2-transduced cells show the mixture of several different sequences, while other cells show only a single sequence. **C. S**equence results of the R750W/sgGata2-transduced cells were submitted to the Synthego website (https://www.synthego.com/), showing various indels in the *Gata2* gene.

**Supplementary figure 7: Peripheral blood analysis of MECOM^WT/R750W^ mice.**

**A.** White blood cells (WBC), hemoglobin (Hb), and platelets (PLT) were measured at 3, 6, 9, and 17 months of age. Data are presented as mean with SD (WT: n = 9, MECOM^WT/R751W^ mice: n = 13).

## Notes

### Competing Interest Statement

The authors have declared no competing interest.

## References

1. Eppert K, Takenaka K, Lechman ER, Waldron L, Nilsson B, van Galen P, et al. Stem cell gene expression programs influence clinical outcome in human leukemia. Nature Medicine 2011 Sep; 17(9): 1086–U1091.

2. Goyama S, Kurokawa M. Pathogenetic significance of ecotropic viral integration site-1 in hematological malignancies. Cancer Science 2009 Jun; 100(6): 990–995.

3. Goyama S, Yamamoto G, Shimabe M, Sato T, Ichikawa M, Ogawa S, et al. Evi-1 is a critical regulator for hematopoietic stem cells and transformed leukemic cells. Cell Stem Cell 2008 Aug; 3(2): 207–220.

4. Kataoka K, Sato T, Yoshimi A, Goyama S, Tsuruta T, Kobayashi H, et al. Evi1 is essential for hematopoietic stem cell self-renewal, and its expression marks hematopoietic cells with long-term multilineage repopulating activity. Journal of Experimental Medicine 2011 Nov; 208(12): 2402–2415.

5. Yuasa H, Oike Y, Iwama A, Nishikata I, Sugiyama D, Perkins A, et al. Oncogenic transcription factor Evi1 regulates hematopoietic stem cell proliferation through GATA-2 expression. Embo Journal 2005 Jun; 24(11): 1976–1987.

6. Morishita K, Parganas E, Willman CL, Whittaker MH, Drabkin H, Oval J, et al. ACTIVATION OF EVI1 GENE-EXPRESSION IN HUMAN ACUTE MYELOGENOUS LEUKEMIAS BY TRANSLOCATIONS SPANNING 300-400 KILOBASES ON CHROMOSOME BAND-3Q26. Proceedings of the National Academy of Sciences of the United States of America 1992 May; 89(9): 3937–3941.

7. Kataoka K, Kurokawa M. Ecotropic viral integration site 1, stem cell self-renewal and leukemogenesis. Cancer Science 2012 Aug; 103(8): 1371–1377.

8. Cui W, Sun JL, Cotta CV, Medeiros LJ, Lin P. Myelodysplastic Syndrome With inv(3)(q21q26.2) or t(3;3)(q21;q26.2) Has a High Risk for Progression to Acute Myeloid Leukemia. American Journal of Clinical Pathology 2011 Aug; 136(2): 282–288.

9. Lugthart S, Groschel S, Beverloo HB, Kayser S, Valk PJM, van Zelderen-Bhola SL, et al. Clinical, Molecular, and Prognostic Significance of WHO Type inv(3)(q21q26.2)/t(3;3)(q21;q26.2) and Various Other 3q Abnormalities in Acute Myeloid Leukemia. Journal of Clinical Oncology 2010 Aug; 28(24): 3890–3898.

10. Rogers HJ, Vardiman JW, Anastasi J, Raca G, Savage NM, Cherry AM, et al. Complex or monosomal karyotype and not blast percentage is associated with poor survival in acute myeloid leukemia and myelodysplastic syndrome patients with inv(3)(q21q26.2)/t(3;3)(q21;q26.2): a Bone Marrow Pathology Group study. Haematologica 2014 May; 99(5): 821–829.

11. Delwel R, Funabiki T, Kreider BL, Morishita K, Ihle JN. 4 OF THE 7 ZINC FINGERS OF THE EVI-1 MYELOID-TRANSFORMING GENE ARE REQUIRED FOR SEQUENCE-SPECIFIC BINDING TO GA(C/T)AAGA(T/C)AAGATAA. Molecular and Cellular Biology 1993 Jul; 13(7): 4291–4300.

12. Perkins AS, Fishel R, Jenkins NA, Copeland NG. Evi-1, a murine zinc finger proto-oncogene, encodes a sequence-specific DNA-binding protein. Molecular and cellular biology 1991; 11(5): 2665–2674.

13. Funabiki T, Kreider BL, Ihle JN. THE CARBOXYL DOMAIN OF ZINC FINGERS OF THE EVI-1 MYELOID TRANSFORMING GENE BINDS A CONSENSUS SEQUENCE OF GAAGATGAG. Oncogene 1994 Jun; 9(6): 1575–1581.

14. Shimabe M, Goyama S, Watanabe-Okochi N, Yoshimi A, Ichikawa M, Imai Y, et al. Pbx1 is a downstream target of Evi-1 in hematopoietic stem/progenitors and leukemic cells. Oncogene 2009 Dec 10; 28(49): 4364–4374.

15. Chiba A, Masamoto Y, Mizuno H, Kurokawa M. GFI1 Is a Downstream Target of EVI1 in Normal Hematopoiesis. Blood 2020 Nov; 136.

16. Masamoto Y, Chiba A, Mizuno H, Hino T, Hayashida H, Sato T, et al. EVI1 exerts distinct roles in AML via ERG and cyclin D1 promoting a chemoresistant and immune-suppressive environment. Blood Advances 2023 Apr; 7(8): 1577–1593.

17. Schmoellerl J, Barbosa IAM, Minnich M, Andersch F, Smeenk L, Havermans M, et al. EVI1 drives leukemogenesis through aberrant ERG activation. Blood 2023 Feb; 141(5): 453–466.

18. Yoshimi A, Goyama S, Watanabe-Okochi N, Yoshiki Y, Nannya Y, Nitta E, et al. Evi1 represses PTEN expression and activates PI3K/AKT/mTOR via interactions with polycomb proteins. Blood 2011 Mar; 117(13): 3617–3628.

19. Izutsu K, Kurokawa M, Imai Y, Maki K, Mitani K, Hirai H. The corepressor CtBP interacts with Evi-1 to repress transforming growth factor beta signaling. Blood 2001 May 1; 97(9): 2815–2822.

20. Palmer S, Brouillet JP, Kilbey A, Fulton R, Walker M, Crossley M, et al. Evi-1 transforming and repressor activities are mediated by CtBP Co-repressor proteins. Journal of Biological Chemistry 2001 Jul 13; 276(28): 25834–25840.

21. Cattaneo F, Nucifora G. EVI1 recruits the histone methyltransferase SUV39H1 for transcription repression. Journal of Cellular Biochemistry 2008 Oct; 105(2): 344–352.

22. Spensberger D, Delwel R. A novel interaction between the proto-oncogene Evi1 and histone methyltransferases, SUV39H1 and G9a. Febs Letters 2008 Aug; 582(18): 2761–2767.

23. Goyama S, Nitta E, Yoshino T, Kako S, Watanabe-Okochi N, Shimabe M, et al. EVI-1 interacts with histone methyltransferases SUV39H1 and G9a for transcriptional repression and bone marrow immortalization. Leukemia 2010 Jan; 24(1): 81–88.

24. de Pater E, Kaimakis P, Vink CS, Yokomizo T, Yamada-Inagawa T, van der Linden R, et al. Gata2 is required for HSC generation and survival. Journal of Experimental Medicine 2013 Dec 16; 210(13): 2843–2850.

25. Ling KW, Ottersbach K, van Hamburg JP, Oziemlak A, Tsai FY, Orkin SH, et al. GATA-2 plays two functionally distinct roles during the ontogeny of hematopoietic stem cells. Journal of Experimental Medicine 2004 Oct; 200(7): 871–882.

26. Ikonomi P, Rivera CE, Riordan M, Washington G, Schechter AN, Noguchi CT. Overexpression of GATA-2 inhibits erythroid and promotes megakaryocyte differentiation. Experimental Hematology 2000 Dec; 28(12): 1423–1431.

27. Huang Z, Dore LC, Li Z, Orkin SH, Feng G, Lin S, et al. GATA-2 Reinforces Megakaryocyte Development in the Absence of GATA-1. Molecular and Cellular Biology 2009 Sep; 29(18): 5168–5180.

28. Ohmori Sy, Moriguchi T, Noguchi Y, Ikeda M, Kobayashi K, Tomaru N, et al. GATA2 is critical for the maintenance of cellular identity in differentiated mast cells derived from mouse bone marrow. Blood 2015 May 21; 125(21): 3306–3315.

29. Li Y, Gao J, Kamran M, Harmacek L, Danhorn T, Leach SM, et al. GATA2 regulates mast cell identity and responsiveness to antigenic stimulation by promoting chromatin remodeling at super-enhancers. Nature Communications 2021 Jan 21; 12(1).

30. Sato T, Goyama S, Nitta E, Takeshita M, Yoshimi M, Nakagawa M, et al. Evi-1 promotes para-aortic splanchnopleural hematopoiesis through up-regulation of GATA-2 and repression of TGF-β signaling. Cancer Science 2008 Jul; 99(7): 1407–1413.

31. Yamazaki H, Suzuki M, Otsuki A, Shimizu R, Bresnick EH, Engel JD, et al. A Remote GATA2 Hematopoietic Enhancer Drives Leukemogenesis in inv(3)(q21;q26) by Activating EVI1 Expression. Cancer Cell 2014 Apr; 25(4): 415–427.

32. Groschel S, Sanders MA, Hoogenboezem R, de Wit E, Bouwman BAM, Erpelinck C, et al. A Single Oncogenic Enhancer Rearrangement Causes Concomitant EVI1 and GATA2 Deregulation in Leukemia. Cell 2014 Apr; 157(2): 369–381.

33. Yamaoka A, Suzuki M, Katayama S, Orihara D, Engel JD, Yamamoto M. EVI1 and GATA2 misexpression induced by inv(3)(q21q26) contribute to megakaryocyte-lineage skewing and leukemogenesis. Blood Advances 2020 Apr 28; 4(8): 1722–1736.

34. Katayama S, Suzuki M, Yamaoka A, Keleku-Lukwete N, Katsuoka F, Otsuki A, et al. GATA2 haploinsufficiency accelerates EVI1-driven leukemogenesis. Blood 2017 Aug; 130(7): 908–919.

35. Niihori T, Ouchi-Uchiyama M, Sasahara Y, Kaneko T, Hashii Y, Irie M, et al. Mutations in MECOM, Encoding Oncoprotein EVI1, Cause Radioulnar Synostosis with Amegakaryocytic Thrombocytopenia. American Journal of Human Genetics 2015 Dec; 97(6): 848–854.

36. Osumi T, Tsujimoto S, Nakabayashi K, Taniguchi M, Shirai R, Yoshida M, et al. Somatic MECOM mosaicism in a patient with congenital bone marrow failure without a radial abnormality. Pediatric Blood & Cancer 2018 Jun; 65(6).

37. Germeshausen M, Ancliff P, Estrada J, Metzler M, Ponstingl E, Rutschle H, et al. MECOM-associated syndrome: a heterogeneous inherited bone marrow failure syndrome with amegakaryocytic thrombocytopenia. Blood Advances 2018 Mar; 2(6): 586–596.

38. Voit RA, Sankaran VG. MECOM Deficiency: from Bone Marrow Failure to Impaired B-Cell Development. Journal of Clinical Immunology 2023 Aug; 43(6): 1052–1066.

39. Ripperger T, Hofmann W, Koch JC, Shirneshan K, Haase D, Wulf G, et al. MDS1 and EVI1 complex locus (MECOM): a novel candidate gene for hereditary hematological malignancies. Haematologica 2018 Feb; 103(2): E55–E58.

40. Shen F, Yang YJ, Zheng Y, Li PC, Luo ZQ, Fu YY, et al. MECOM-related disorder: Radioulnar synostosis without hematological aberration due to unique variants. Genetics in Medicine 2022 May; 24(5): 1139–1147.

41. Walne A, Tummala H, Ellison A, Cardoso S, Sidhu J, Sciuccati G, et al. Expanding the phenotypic and genetic spectrum of radioulnar synostosis associated hematological disease. Haematologica 2018 Jul; 103(7): E284–E287.

42. Nagai K, Niihori T, Muto A, Hayashi Y, Abe T, Igarashi K, et al. Mecom mutation related to radioulnar synostosis with amegakaryocytic thrombocytopenia reduces HSPCs in mice. Blood advances 2023 2023-09-26; 7(18): 5409–5420.

43. Platt RJ, Chen SD, Zhou Y, Yim MJ, Swiech L, Kempton HR, et al. CRISPR-Cas9 Knockin Mice for Genome Editing and Cancer Modeling. Cell 2014 Oct; 159(2): 440–455.

44. Kitamura T, Koshino Y, Shibata F, Oki T, Nakajima H, Nosaka T, et al. Retrovirus-mediated gene transfer and expression cloning: Powerful tools in functional genornics. Experimental Hematology 2003 Nov; 31(11): 1007–1014.

45. Goyama S, Schibler J, Gasilina A, Shrestha M, Lin S, Link KA, et al. UBASH3B/Sts-1-CBL axis regulates myeloid proliferation in human preleukemia induced by AML1-ETO. Leukemia 2016 Mar; 30(3): 728–739.

46. Kim D, Landmead B, Salzberg SL. HISAT: a fast spliced aligner with low memory requirements. Nature Methods 2015 Apr; 12(4): 357–U121.

47. Robinson MD, McCarthy DJ, Smyth GK. edgeR: a Bioconductor package for differential expression analysis of digital gene expression data. Bioinformatics 2010 Jan; 26(1): 139–140.

48. Langmead B, Trapnell C, Pop M, Salzberg SL. Ultrafast and memory-efficient alignment of short DNA sequences to the human genome. Genome Biology 2009; 10(3).

49. Zhang Y, Liu T, Meyer CA, Eeckhoute J, Johnson DS, Bernstein BE, et al. Model-based Analysis of ChIP-Seq (MACS). Genome Biology 2008; 9(9).

50. Yu GC, Wang LG, He QY. ChIPseeker: an R/Bioconductor package for ChIP peak annotation, comparison and visualization. Bioinformatics 2015 Jul; 31(14): 2382–2383.

51. Ramírez F, Ryan DP, Grüning B, Bhardwaj V, Kilpert F, Richter AS, et al. deepTools2: a next generation web server for deep-sequencing data analysis. Nucleic Acids Research 2016 Jul; 44(W1): W160–W165.

52. Robinson JT, Thorvaldsdóttir H, Winckler W, Guttman M, Lander ES, Getz G, et al. Integrative genomics viewer. Nature Biotechnology 2011 Jan; 29(1): 24–26.

53. Heinz S, Benner C, Spann N, Bertolino E, Lin YC, Laslo P, et al. Simple Combinations of Lineage-Determining Transcription Factors Prime *cis*-Regulatory Elements Required for Macrophage and B Cell Identities. Molecular Cell 2010 May; 38(4): 576–589.

54. Nitta E, Izutsu K, Yamaguchi Y, Imai Y, Ogawa S, Chiba S, et al. Oligomerization of Evi-1 regulated by the PR domain contributes to recruitment of corepressor CtBP. Oncogene 2005 Sep; 24(40): 6165–6173.

55. Hsu AP, Sampaio EP, Khan J, Calvo KR, Lemieux JE, Patel SY, et al. Mutations in GATA2 are associated with the autosomal dominant and sporadic monocytopenia and mycobacterial infection (MonoMAC) syndrome. Blood 2011 Sep 8; 118(10): 2653–2655.

56. Hoyt PR, Bartholomew C, Davis AJ, Yutzey K, Gamer LW, Potter SS, et al. The Evil proto-oncogene is required at midgestation for neural, heart, and paraxial mesenchyme development. Mechanisms of Development 1997 Jul; 65(1-2): 55–70.

