## Supplementary Figure1-7 for "MECOM promotes leukemia progression and inhibits mast cell differentiation through functional competition with GATA2"

### Supplementary Figure 1

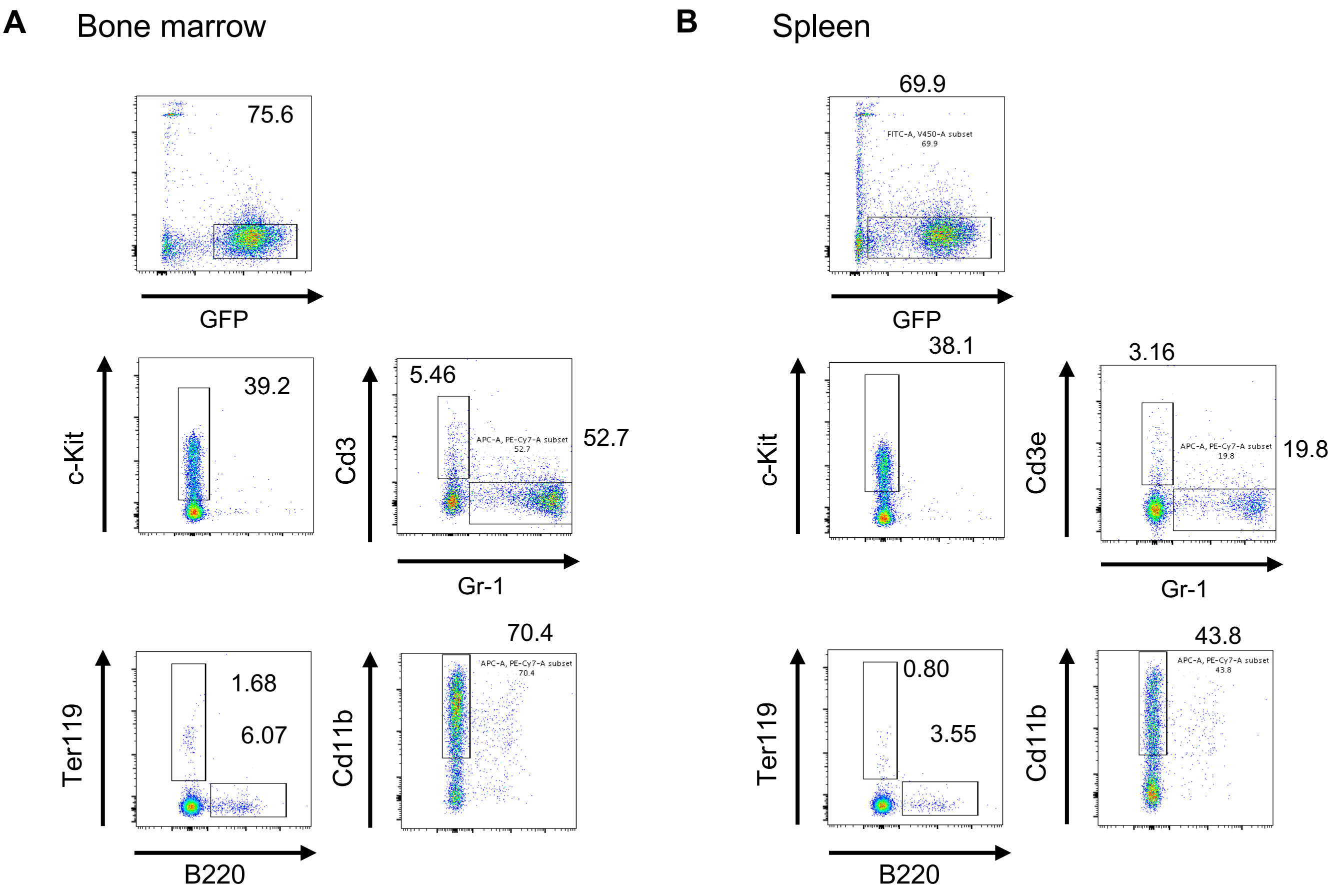

### Supplementary Figure 2

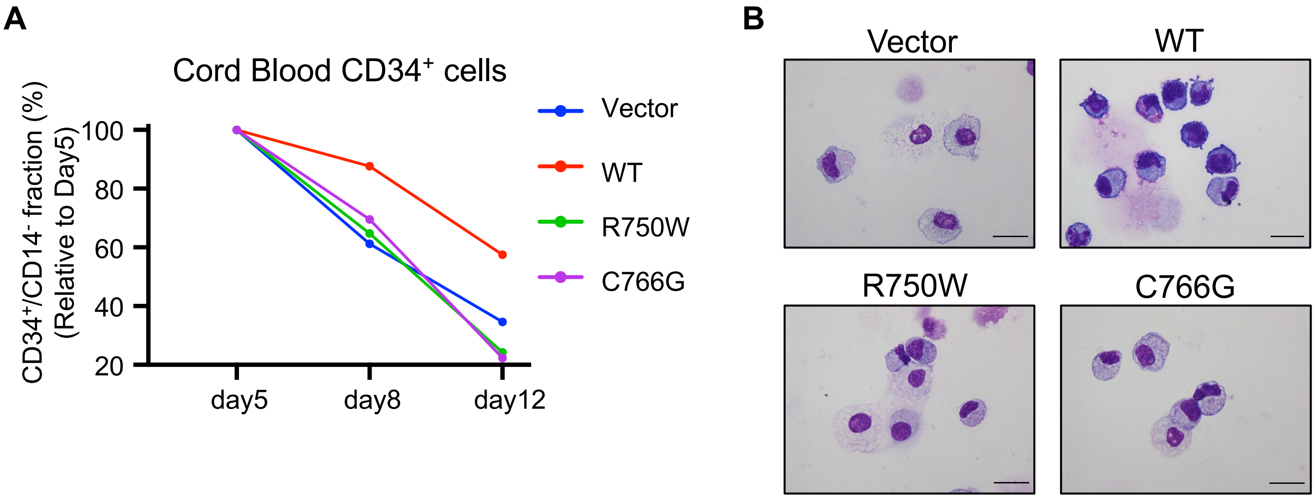

### Supplementary Figure 3

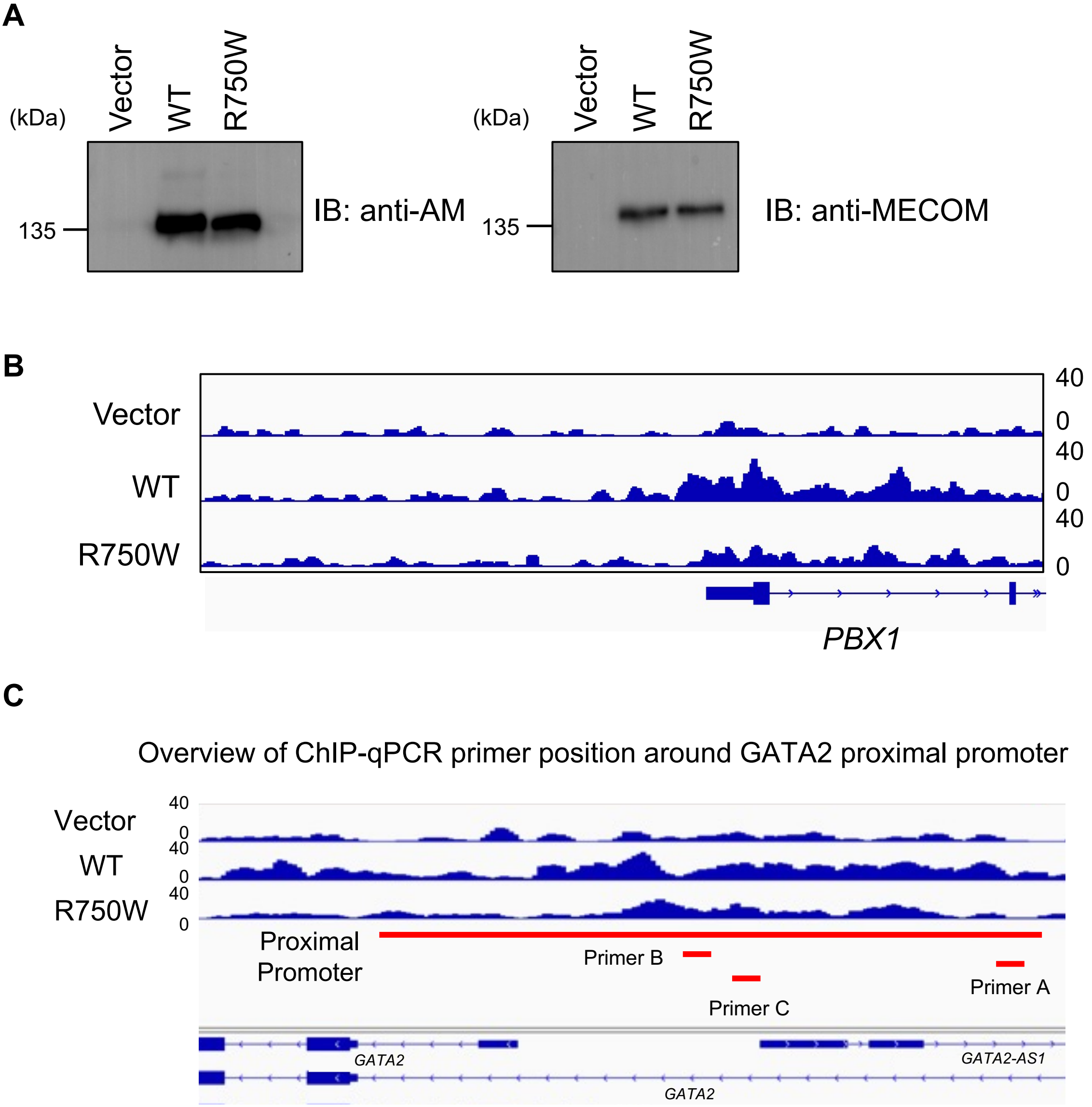

### Supplementary Figure 4

**A**

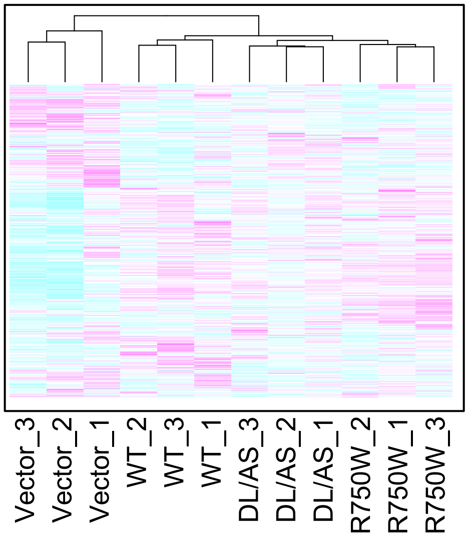


**B**

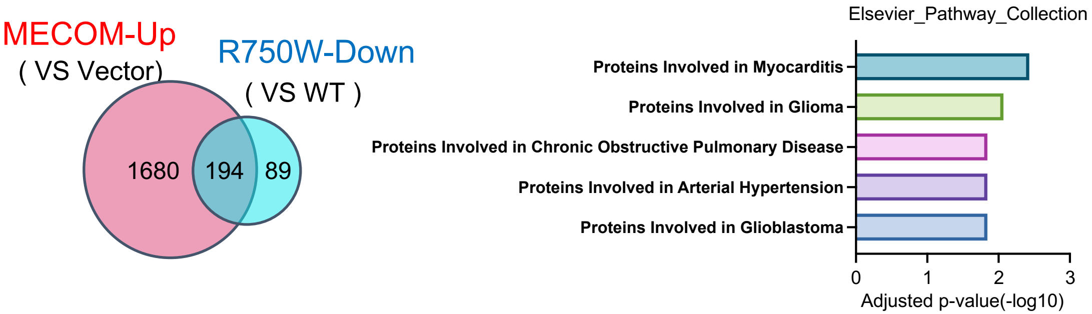


**C**

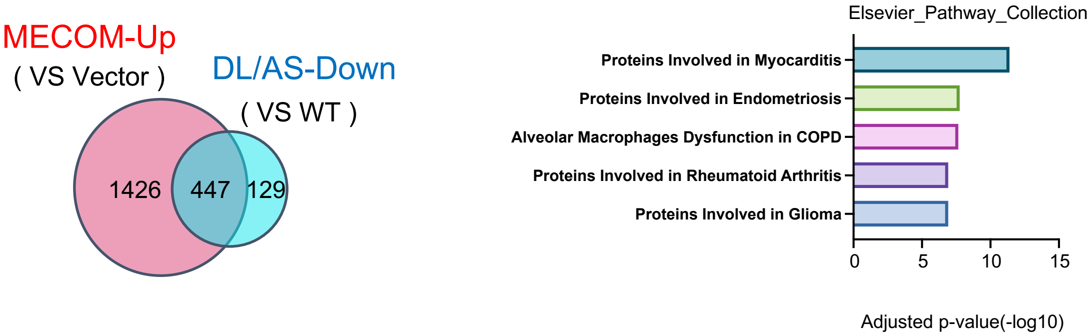

### Supplementary Figure 5

**A**

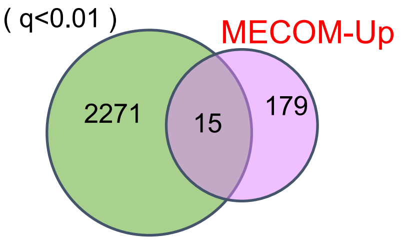


| | | |
|---|---|---|
| ARID5B | FSCN1 | SEMA4A |
| ABTB2 | SERPINB6 | COL6A1 |
| TRIB1 | BHLHE40 | KITLG |
| OSBPL6 | PLXNB2 | |
| PLOD2 | PCDH9 | |
| RNF213 | SOCS2 | |

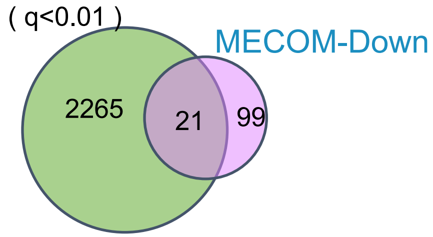


| | | |
|---|---|---|
| SOS1 | HMGN4 | DUSP10 |
| EPB41L4A | ARL4C | ZNF521 |
| MITF | LMO4 | KIAA0040 |
| NID1 | CNRIP1 | OGFRL1 |
| PDCD4 | ELF1 | NEO1 |
| CDC42BPA | CCND1 | ADAMTS3 |
| KCNH2 | NDFIP2 | EPAS1 |

**B**

Overview of ChIP-qPCR primer position around MITF promoter

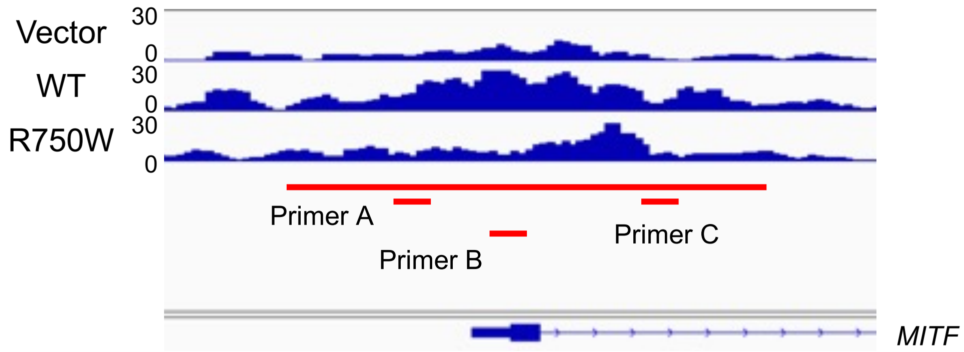


**C**

MITF expression in mouse BM cells

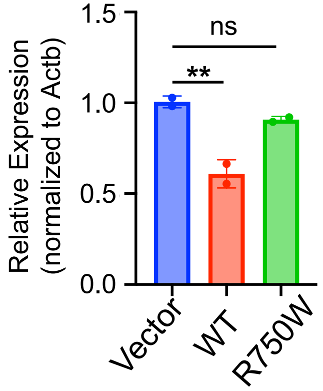

### Supplementary Figure 6

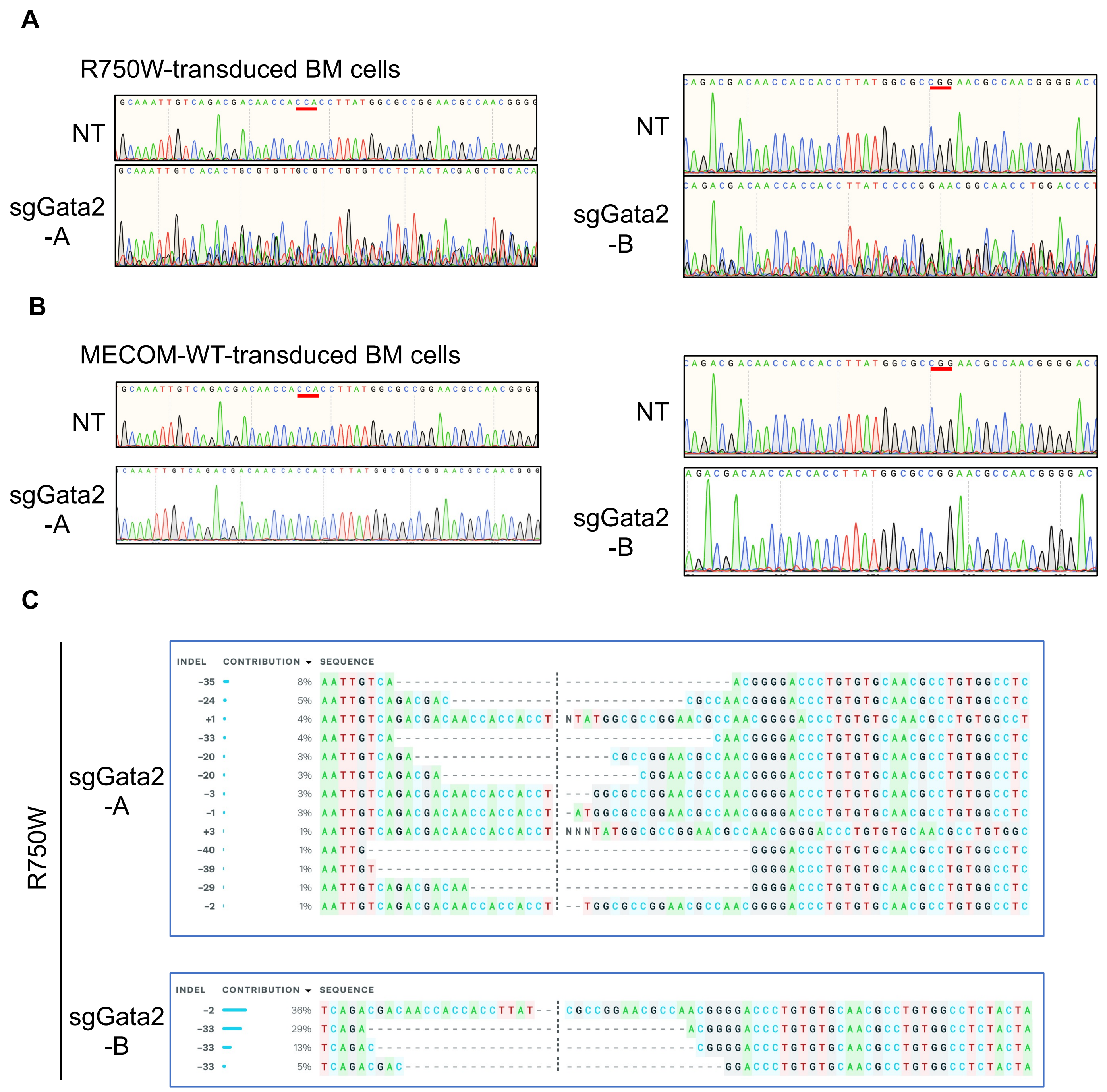

### Supplementary Figure 7

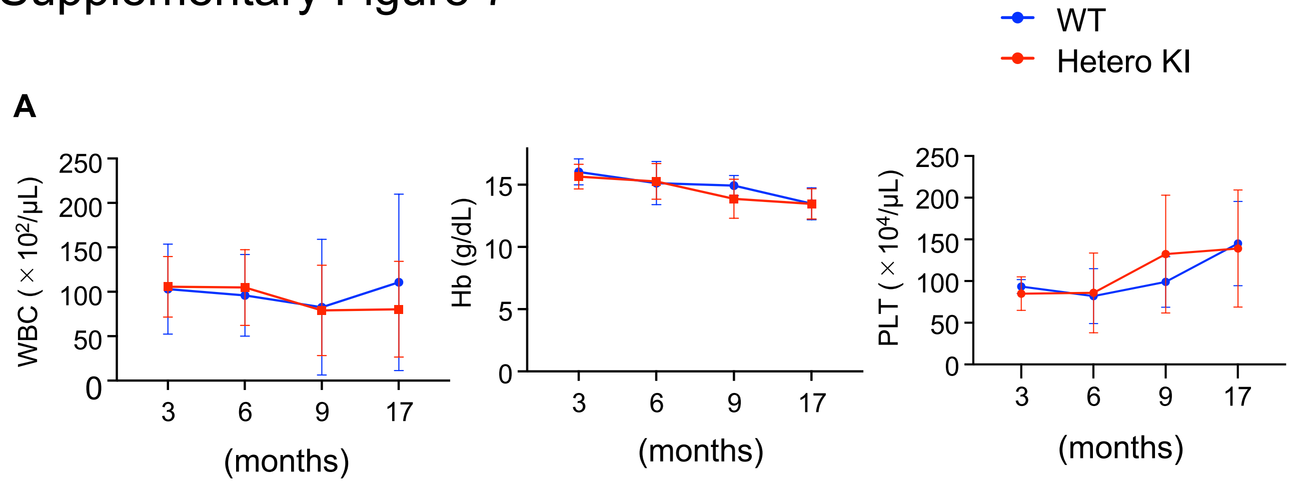
